# Structural analysis of the PTEN:P-Rex2 signalling node reveals how cancer-associated mutations coordinate to hyperactivate Rac1

**DOI:** 10.1101/2020.04.11.036434

**Authors:** Laura D’Andrea, Christina M. Lucato, Elsa A. Marquez, Yong-Gang Chang, Srgjan Civciristov, Cheng Huang, Hans Elmlund, Ralf B. Schittenhelm, Christina A. Mitchell, James C. Whisstock, Michelle L. Halls, Andrew M. Ellisdon

**Author notes:** These authors contributed equally (Laura D’Andrea, Christina M. Lucato). Correspondence should be addressed to A.M.E. or M.L.H.

## Abstract

The PTEN:P-Rex2 complex is one of the most commonly mutated signaling nodes in metastatic cancer. Assembly of the PTEN:P-Rex2 complex inhibits the activity of both proteins, and its dysregulation can drive PI3K-AKT signaling and cell proliferation. Here, using extensive crosslinking mass spectrometry and functional studies, we provide crucial mechanistic insights into PTEN:P-Rex2 complex assembly and co-inhibition. PTEN is anchored to P-Rex2 by interactions between the PTEN PDZ-BM tail and the second PDZ domain of P-Rex2. This interaction bridges PTEN across the P-Rex2 surface, occluding PTEN membrane-binding and PI(3,4,5)P_3_ hydrolysis. Conversely, PTEN both allosterically promotes an autoinhibited P-Rex2 conformation and occludes Gβγ binding and GPCR activation. These insights allow us to define a new gain-of-function class of cancer mutations within the PTEN:P-Rex2 interface that uncouples PTEN inhibition of Rac1 signaling. These findings provide a mechanistic framework to understand the dysregulation of the PTEN:P-Rex2 signaling node in metastatic cancer.

## INTRODUCTION

Phosphatase and tensin homologue (PTEN) is a dual-specificity phosphatase with a canonical tumour suppressor role in converting the lipid second messenger phosphatidylinositol (3,4,5)-Tris-HCl-phosphate (PI(3,4,5)P_3_) to PI(4,5)P_2_ to inhibit PI3K-AKT signaling^1^. In addition to its phosphatase-dependent roles, PTEN can act as a tumour suppressor by providing a scaffold to bind and regulate multiple proteins and protein complexes^2^. For example, PTEN forms a crucial co-inhibitory complex with the PI(3,4,5)P_3_-dependent Rac-exchange factor 2 (P-Rex2) guanine nucleotide exchange factor (GEF) that catalyses nucleotide exchange for the Rac-like RhoGTPase subfamily^3^. Assembly of the PTEN:P-Rex2 complex results in enhanced PI3K-AKT signaling and cell proliferation^3^, and suppression of Rac1-mediated cell migration and invasion^4^ (Supplementary Fig. 1a). P-Rex2 is autoinhibited in the basal state and activated at the membrane upon binding PI(3,4,5)P_3_ and Gβγ downstream of receptor tyrosine kinase (RTK) and G protein-coupled receptor (GPCR) stimulation. As such, in addition to its PI(3,4,5)P_3_-independent scaffolding role in P-Rex2 inhibition, PTEN can also indirectly inhibit P-Rex2 in a phosphatase-dependent manner by reducing PI(3,4,5)P_3_ levels at the membrane.

PTEN comprises an N-terminal dual-specificity phosphatase domain (DUSP), a C2 domain, and an unstructured carboxy-terminal tail that ends in a PDZ-binding motif (PDZ-BM) (Fig. 1a)^5,6^. The DUSP domain contains the CX_5_R motif common to protein tyrosine phosphatases and is sandwiched against a C2 domain that regulates membrane association^5,6^. PTEN is flexible, alternating between open and closed conformations upon PTEN-tail phosphorylation and subsequent binding of the PTEN-tail back to inhibit the globular DUSP and C2 domains^5,7,8^.

**Fig. 1.**
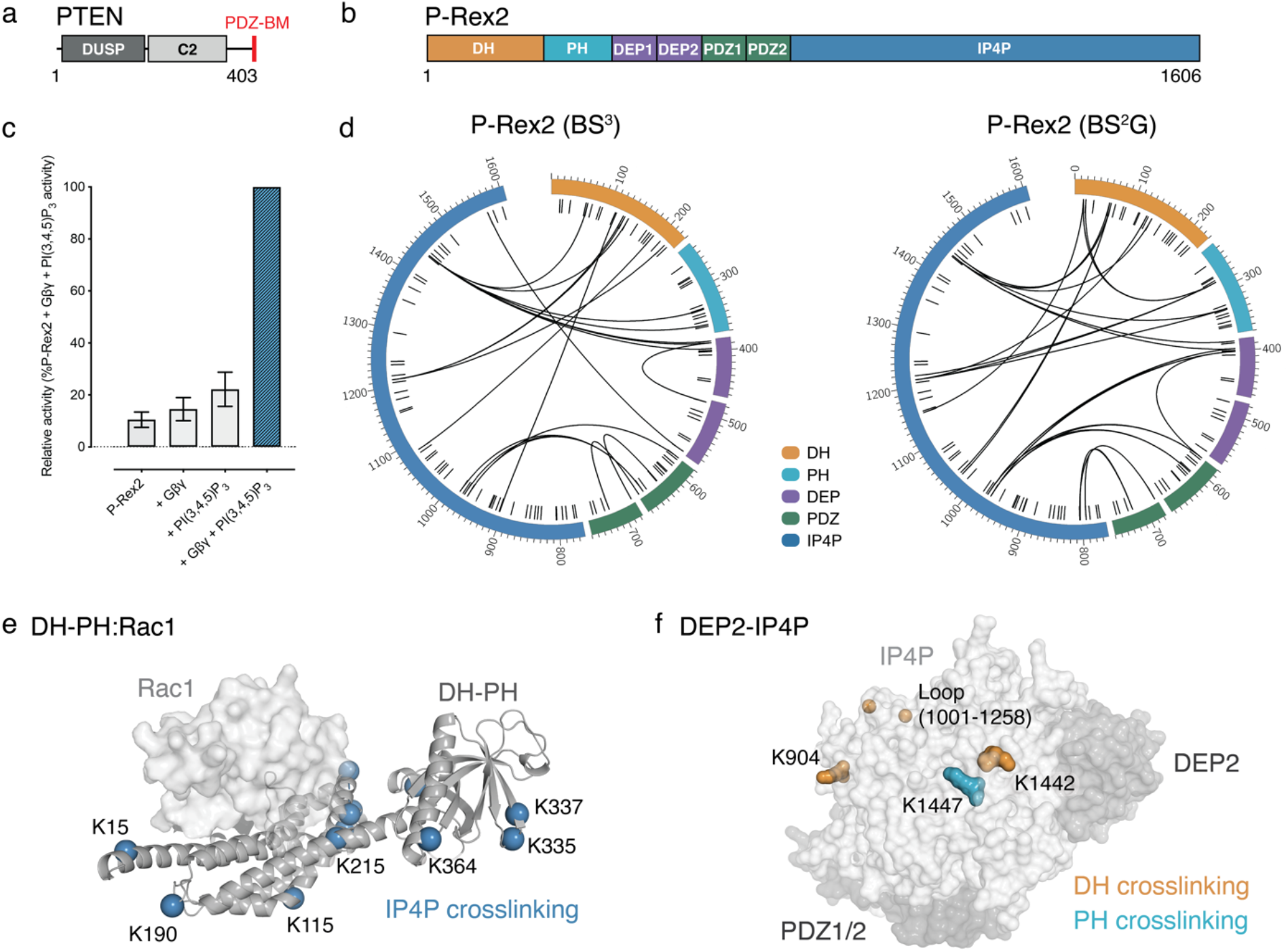
P-Rex2 is locked in a closed conformation by the IP4P domain. **a**, Domain layout of PTEN. **b**, Domain layout of P-Rex2. **c**, Rac1 ^35^S-GTPγS loading by P-Rex2 in the presence or absence of Gβγ, PI(3,4,5)P_3_-containing liposomes, or combined Gβγ and PI(3,4,5)P_3_-containing liposomes. **d**, P-Rex2 intramolecular BS^3^ and BS^2^G crosslinks identified by CLMS. Black lines show the identified crosslinks and P-Rex2 domains are coloured as per **b**. Short black dashes correspond to lysine residues within the primary sequence. Intradomain crosslinks are omitted for clarity. **e**, Intramolecular crosslinking sites mapped to the P-Rex2 DH-PH domains (modeled from 4YON^10^). **f**, Interdomain crosslinking sites mapped to the P-Rex2 DEP2-IP4P domains (modeled from 6PCV^12^).

Similarly, P-Rex2 alternates between an autoinhibited basal state and an activated membrane-associated conformation. P-Rex2 contains an N-terminal DH-PH tandem domain that forms the minimal RhoGEF catalytic subunit^3^. C-terminal to the DH-PH domains are two DEP domains, two PDZ domains, and a 4-phosphatase homology domain (IP4P) (Fig. 1b). There are no reported structures of P-Rex2 outside of the isolated PH domain^9^. However, the DH-PH domains of the P-Rex2 homologue P-Rex1 resemble the typical RhoGEF structure and catalytic mechanism^10,11^. More recently, the P-Rex1 DEP2-IP4P:Gβγ complex structure has revealed the organisation of the C-terminal tail of P-Rex1, with a central IP4P domain flanked by the DEP2 and tandem PDZ domains with Gβγ binding to the surface of the IP4P domain^12^. However, the orientation of the catalytic DH-PH domains to the autoinhibitory C-terminal domains is yet to be resolved. As such, there remains minimal insight into the mechanism of P-Rex family autoinhibition and activation upon PI(3,4,5)P_3_ and Gβγ binding.

PTEN:P-Rex2 complex assembly occurs independently of both PTEN phosphatase activity^13^ and P-Rex2 GEF activity^14^. However, the flexibility of both PTEN and P-Rex2 has limited high-resolution insights into the assembly of the complex. Domain deletion studies suggest that the P-Rex2 DH-PH domains are necessary to inhibit PTEN activity, mediated by an interaction between the P-Rex2 PH domain and the PTEN DUSP and C2 domains^14^. A second binding site between the PTEN PDZ-BM and the P-Rex2 IP4P domain was sufficient to suppress P-Rex2 GEF-mediated breast cancer cell invasion^4^. However, studies diverge on the requirement for the PDZ-BM for complex assembly^3^.

P-Rex2 is mutated frequently in cancer. In melanoma, the incidence of P-Rex2 mutations has been reported as high as 27% within the TCGA cohort and 14% in a separate cohort of 107 melanomas^15–17^. High-incidence of P-Rex2 mutations has also been reported in pancreatic, breast, and lung cancers^18–20^. More broadly, P-Rex2 is the fourth most commonly mutated oncogene in the MET500 cohort of metastatic cancer, underlining a potential role as a metastatic driver^21^. Strikingly, within the same MET500 cohort, PTEN is the third most commonly mutated tumour-suppressor protein^21^. This combined incidence highlights the PTEN:P-Rex2 complex as one of the most commonly mutated signaling nodes in metastatic cancer.

PTEN cancer-associated mutations are generally clustered to the catalytic domain (e.g. C124S) and frequently result in PTEN suppression and enhanced PI3K-AKT signaling^5^. P-Rex2 cancer-associated somatic mutations are not clustered in hot-spots, but range throughout the primary sequence^15,19,21^. As we know little about the structure of P-Rex2 or its interaction with PTEN, it has proven difficult to predict the importance of individual mutations to the dysregulation of PI3K-AKT and RhoGTPase signaling. For example, melanoma-associated missense mutations have proven challenging to predict in terms of functional impact with some (e.g. G884D) accelerating in vivo tumorigenesis while others (e.g. P948S) do not^15^. P-Rex2 missense mutations also display differential abilities to avoid PTEN inhibition of their GEF activity^4^. P-Rex2 truncation mutations can accelerate tumorigenesis when ectopically expressed in melanocytes transplanted into immunodeficient mice^15^ while also increasing tumour penetrance^13^. P-Rex2 truncation mutations are thought to decrease P-Rex2 autoinhibition while avoiding PTEN binding and inhibition^13^. P-Rex2 mutations are often associated with high PTEN expression^4^, although it is unclear if there is a synergy between PTEN and P-Rex2 cancer-associated mutations.

Here, we use a combination of crosslinking mass spectrometry (CLMS) and extensive functional studies to build a mechanistic framework to understand PTEN:P-Rex2 complex assembly, co-inhibition, and dysregulation in cancer. We show that the P-Rex2 PDZ2 and IP4P domains form the essential PTEN binding site. PTEN binding inhibits P-Rex2 by both blocking Gβγ activation and allosterically supporting an autoinhibited P-Rex2 conformation. Conversely, PTEN binds across the P-Rex2 surface, occluding PI(3,4,5)P_3_ hydrolysis by PTEN. Functional studies define a new class of gain-of-function P-Rex2 cancer mutations situated within the PTEN:P-Rex2 interface, which prevent the assembly of the co-inhibitory complex and activate Rac1 signaling. Finally, we demonstrate synergy between PTEN deactivating and P-Rex2 truncation mutations that combine to drive Rac1 activation to a greater extent than either single mutation alone.

## RESULTS

### The P-Rex2 IP4P domain locks P-Rex2 in a closed autoinhibited conformation

In the basal state, the P-Rex2 catalytic N-terminal DH-PH tandem domains are autoinhibited by an unknown mechanism that requires the C-terminal domains (Fig. 1b)^13^. In agreement, we observed that recombinant P-Rex2 (Supplementary Fig. 1b) was autoinhibited and required the addition of both Gβγ and PI(3,4,5)P_3_ to efficiently exchange GDP for GTP on Rac1 (Fig. 1c). We were unable to detect binding between full-length P-Rex2 and Rac1 by surface plasmon resonance (SPR), compared to the high-affinity interaction observed between the isolated P-Rex2 DH-PH domains and Rac1 (Supplementary Fig. 1c). These data suggest that autoinhibition is caused by steric occlusion of the Rac1 binding site on the DH domain by the C-terminal domains of P-Rex2.

CLMS is a powerful technique that provides amino-acid proximity measurements of native proteins in solution. This technique has enabled many recent insights into the molecular determinants of protein-protein interactions and the mapping of subunits within large protein complexes^22^. We, therefore, used CLMS to gain insight into the mechanism of P-Rex2 autoinhibition. BS^3^ (11.4 Å linker length) and BS^2^G (7.7 Å linker length) lysine crosslinkers identified a total of 49 unique interdomain crosslinks (20 BS^3^ and 29 BS^2^G) within autoinhibited P-Rex2 (Fig. 1d and Supplementary Table 1). Validation of our CLMS approach revealed that 94% of all crosslinks mapped to allowable regions of P-Rex2 models (<30 Å between Cα atoms^23^) (Supplementary Fig. 1d). Interestingly, we consistently observed incompatible long-range crosslinks between the P-Rex2 DH and PH domains suggesting a different arrangement of these domains in autoinhibited P-Rex2 compared to the isolated and active P-Rex1 DH-PH:Rac1 complex used for structural modeling^10^ (Supplementary Fig. 1d).

The extent of the interdomain crosslinks reveals a closed globular structure, rather than an extended conformation for autoinhibited P-Rex2. The IP4P domain is central to this closed conformation, as it crosslinks to each of the six remaining P-Rex2 domains (Fig. 1d). In particular, the IP4P domain forms numerous contacts with the catalytic DH-PH domains and the PH-DEP1 linker region. In total, we identified 12 unique sites in the DH-PH domains, including multiple lysine residues that occur in the modeled DH-PH:Rac1 interface (Fig. 1d,e).

The crosslinks between the IP4P and DH-PH domains are initiated from ten IP4P sites across a surface formed by helical and loop regions (residues 1001-1258) of the IP4P domain distal to the PDZ and DEP domains (Fig. 1d,f). The loop region of the IP4P domain is predominantly unstructured in cryo-EM studies of the homologous P-Rex1 DEP2-IP4P regions^12^. These data support a model of P-Rex2 autoinhibition where the IP4P domain is fundamental to maintaining an inactive, closed conformation by embracing the DH-PH domains to prevent GTPase binding.

### Dual inhibition of PTEN and P-Rex2: the PTEN:P-Rex2 interface occludes the PTEN active site and promotes a closed P-Rex2 conformation

To examine the structural basis for PTEN and P-Rex2 co-inhibition, we expanded our CLMS studies to the PTEN:P-Rex2 complex. As per previous studies, we confirmed that PTEN:P-Rex2 complex formation co-inhibited both P-Rex2 GEF and PTEN phosphatase activity (Fig. 2a,b). Next, we assembled a stoichiometric PTEN:P-Rex2 complex by size-exclusion chromatography (Fig. 2c and Supplementary Fig. 2a) and performed CLMS of the PTEN:P-Rex2 complex (Fig. 2d,e, Supplementary Fig. 2b,c and Supplementary Table 1). Validation mapping showed that all crosslink sites in PTEN were within the expected distance constraints, and 91% of the P-Rex2 crosslink sites were in allowable regions (Supplementary Fig. 2d).

**Fig. 2.**
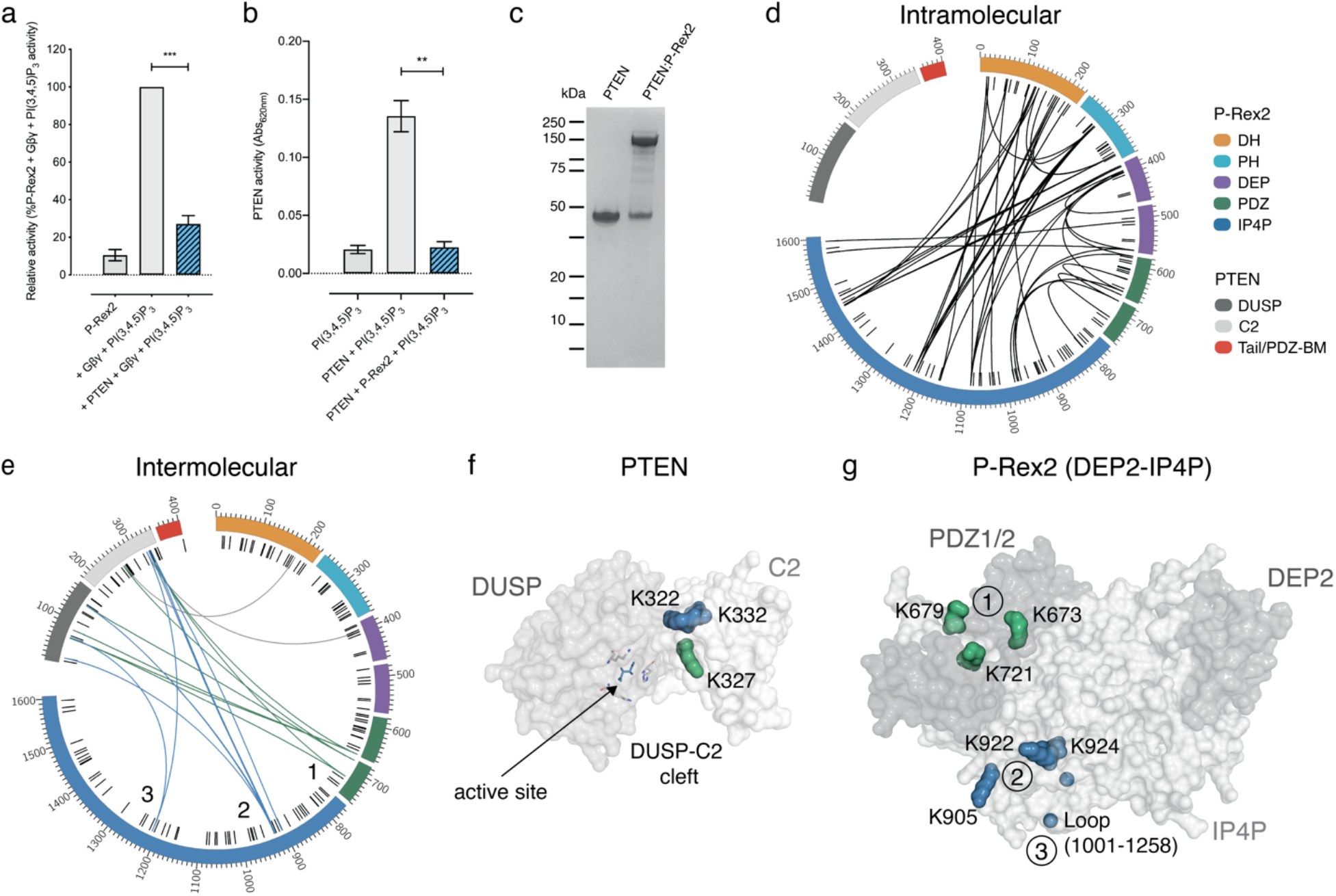
PTEN binds to the P-Rex2 PDZ2 and IP4P domains. **a**, Rac1 ^35^S-GTPγS loading by P-Rex2, or the PTEN:P-Rex2 complex, in the presence of Gβγ and PI(3,4,5)P_3_-containing liposomes. **b**, Malachite-green assay of free-phosphate release from PI(3,4,5)P_3_ upon incubation with PTEN or the PTEN:P-Rex2 complex. **c**, SDS-PAGE of insect-cell purified PTEN and the PTEN:P-Rex2 complex. **d**, P-Rex2 intramolecular BS^3^ crosslinks within the PTEN:P-Rex2 complex. **e**, Intermolecular BS^3^ crosslinks identified within the PTEN:P-Rex2 complex. **f**, PTEN structure (PDB 1D5R^6^) with PTEN Cα2 lysine residues that interact with the P-Rex2 PDZ2 domain in green and P-Rex2 IP4P domain in blue. PTEN catalytic residues are displayed as sticks. **g**, P-Rex2 DEP2-IP4P domains (modeled from 6PCV^12^) with PDZ domain crosslinks mapped in green and IP4P crosslinks mapped in blue. The three P-Rex2 interacting regions in **e** are highlighted.

Overall, we observed a marked increase in the number of residues forming intramolecular P-Rex2 crosslinks, characterised by a greater distribution of crosslinks between the IP4P domain and the catalytic DH-PH domains (Fig. 2d, Supplementary Fig. 2e and Supplementary Table 1). However, there is no major change in the pattern of intramolecular P-Rex2 crosslinks in the PTEN:P-Rex2 complex (Fig. 2d). These data indicate that PTEN binds to the autoinhibited conformation of P-Rex2 without eliciting rearrangement of the P-Rex2 domain topology. Overall, the increase in crosslinking sites is consistent with PTEN binding and promoting an autoinhibited and closed P-Rex2 conformation.

We then analysed the intermolecular crosslinks between PTEN and P-Rex2. In total, we identified 44 intermolecular crosslinks between PTEN and P-Rex2 corresponding to 18 unique sites within the PTEN:P-Rex2 complex (Fig. 2e and Supplementary Table 1). The crosslinks from P-Rex2 map to the catalytic cleft of the PTEN C2 domain and the surface of the PTEN DUSP domain (Fig. 2e). Three PTEN C2 domain residues (Lys322, Lys327, Lys332), clustered on the Cα2 helix on the surface of the PTEN DUSP-C2 cleft, account for 75% of all observed crosslinks (Fig. 2f). This critical membrane-binding region^24,25^ sits only ~10 Å from the PTEN active site, and therefore, P-Rex2 docking close to this site would sterically occlude PTEN binding to PI(3,4,5)P_3_ and the membrane. To further investigate this possibility, we performed PTEN-liposome pulldown experiments using either PTEN, P-Rex2, or the PTEN:P-Rex2 complex. Given the propensity of the P-Rex2 PH and DEP domains to bind liposomes, we utilised a N-terminally truncated P-Rex2 (PDZ1-IP4P) variant in the pulldown experiments that is unable to bind liposomes but retains PTEN binding. Consistent with a model of steric occlusion, we observed reduced liposome binding to the PTEN:P-Rex2 complex compared to PTEN alone (Supplementary Fig. 2f,g).

Despite previous studies suggesting a central role for the P-Rex2 DH-PH domains in PTEN binding^3,14^, we observed only a few crosslinks between PTEN and the DH (Lys190) and DEP1 (Lys400) domains (Fig. 2e, grey). Instead, we found that crosslinks in the PTEN:P-Rex2 complex arise from three primary locations within P-Rex2: the PDZ2 domain (Site 1; Lys673, Lys679, Lys721) and two regions in the IP4P domain (Site 2; Lys905, Lys922, Lys924 and Site 3; Lys1212) (Fig. 2e,g). Despite a significant distance between the three sites in terms of the primary sequence, all three sites cluster to the same surface of the modeled P-Rex2 structure (Fig. 2g). These data indicate that PTEN binds to P-Rex2 by bridging both the P-Rex2 PDZ2 and IP4P domains.

### PTEN binding blocks Gβγ activation of P-Rex2 downstream of GPCR activation

Comparison between the PTEN:P-Rex2 crosslinking sites, and the P-Rex1:Gβγ cryo-EM structure^12^, indicate that the P-Rex2 PDZ2 domain plays a crucial role in coordinating Gβγ as well as PTEN binding (Fig. 3a). The overlap of the PTEN and Gβγ binding sites in the P-Rex2 PDZ2 domain suggests that PTEN binding may inhibit P-Rex2 activation downstream of GPCR signaling by occluding the Gβγ binding site. To examine this possibility, we first measured binding of P-Rex2, or the PTEN:P-Rex2 complex, to Gβγ by SPR. As expected, P-Rex2 bound Gβγ with an affinity of 100 nM in line with a physiological role of Gβγ in P-Rex2 activation (Fig. 3b). However, we found the PTEN:P-Rex2 complex failed to bind Gβγ across a wide concentration range in agreement with our model of PTEN inhibition of P-Rex2 by steric occlusion of Gβγ binding (Fig. 3b).

**Fig. 3.**
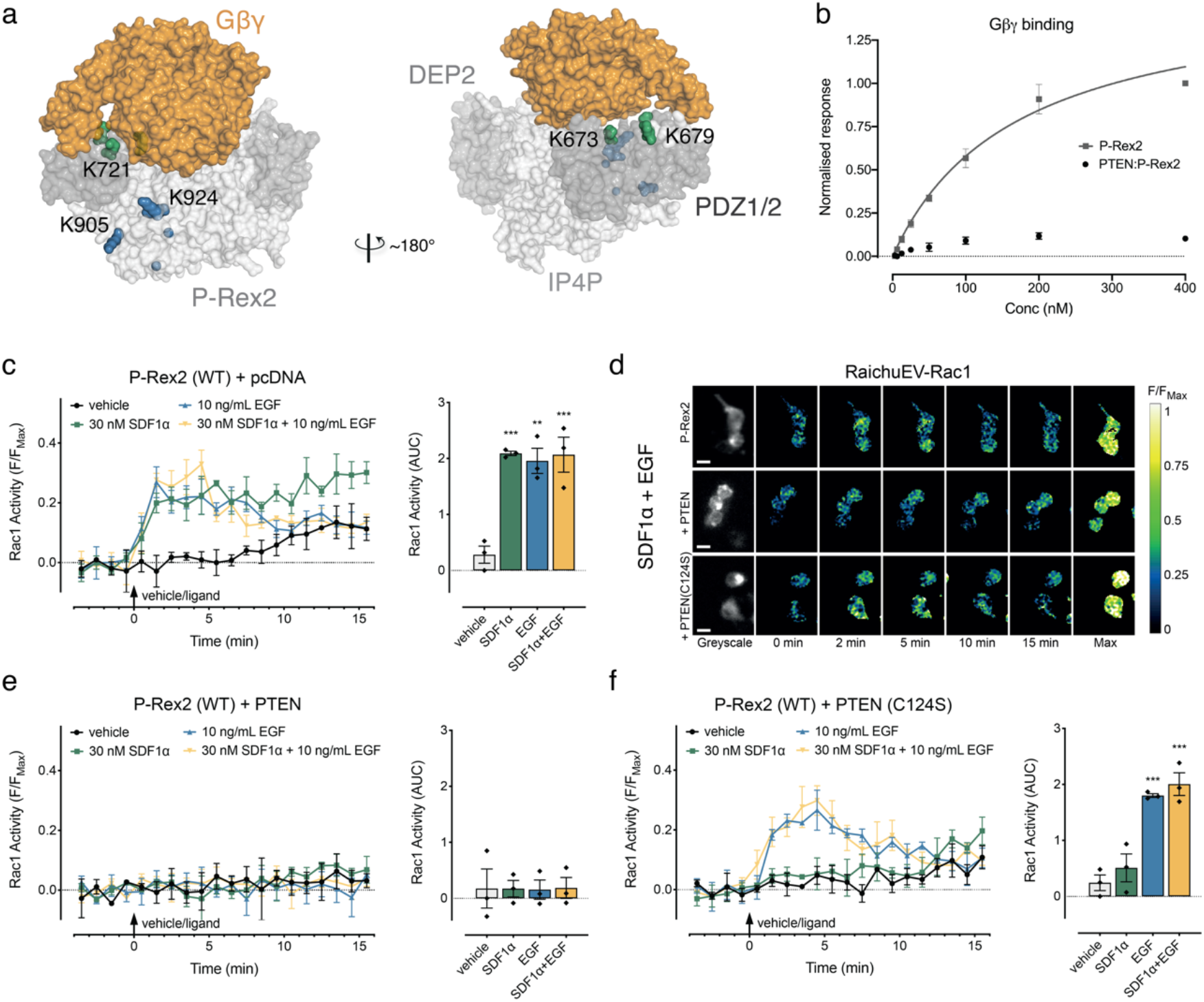
PTEN binding inhibits Gβγ activation of P-Rex2. **a**, Model of P-Rex2 DEP2-IP4P:Gβγ (modeled from 6PCV^12^) highlighting overlap of the Gβγ binding site with PTEN crosslinking sites. **b**, SPR steady-state response curves of Gβγ binding by P-Rex2 and the PTEN:P-Rex2 complex. **c**, Rac1 activity upon transient expression of P-Rex2 (WT) in HEK293 cells. **d**, Representative ratiometric pseudocolour images of HEK293 cells expressing RaichuEV-Rac1 and indicated constructs at baseline (0 min) following stimulation with SDF1α and EGF and the maximal FRET response to the positive control (Max). **e**, Rac1 activity upon transient expression of P-Rex2 (WT) and PTEN (WT) in HEK293 cells. **f**, Rac1 activity upon transient expression of P-Rex2 (WT) and PTEN (C124S) in HEK293 cells. AUC, the area under the curve. Scatter plots show individual data points, symbols/bars represent means, and error bars indicate S.E.M. ** p<0.01 and *** p<0.001 versus vehicle control, one-way ANOVA with Dunnett’s multiple comparison test.

RhoGTPases such as Rac1, RhoA, and Cdc42 are often dysregulated in cancer and contribute to disease progression by promoting migration and metastasis of cancer cells^26^. To investigate PTEN inhibition of P-Rex2 in a cell system, we used HEK293 cells co-transfected with P-Rex2 and the RaichuEV-Rac1 FRET-based biosensor (Supplementary Fig. 3a-c). HEK293 cells have previously been used to monitor P-Rex2 activity and serve as a simple cellular system to delineate the role of the PTEN:P-Rex2 complex in Rac1 activation^4^.

Here, treatment of HEK293 cells with 10 ng/ml EGF (EGFR ligand) caused a rapid and transient increase in Rac1 activity, which declined to baseline by 10 min (Fig. 3c). In contrast, 30 nM SDF1α (CXCR4 ligand) caused a sustained increase in Rac1 activity over the 15 min time course (Fig. 3c). Combined stimulus with both EGF and SDF1α did not reveal any additive effect on Rac1 activation (Fig. 3c,d).

Co-transfection of HEK293 cells with PTEN and P-Rex2 resulted in complete inhibition of Rac1 activity downstream of both EGF and SDF1α stimulus (Fig. 3d,e and Supplementary Fig. 3a-c). As P-Rex2 requires PI(3,4,5)P_3_ for activation, PTEN co-transfection may inhibit P-Rex2 directly by forming a co-inhibitory PTEN:P-Rex2 complex, or indirectly by reducing the available pool of PI(3,4,5)P_3_ at the plasma membrane (Supplementary Fig. 1a). To delineate between these two possibilities, we used a catalytically-dead cancer-associated mutant PTEN (C124S), which is unable to hydrolyse PI(3,4,5)P_3_. In HEK293 cells co-transfected with P-Rex2 and PTEN (C124S), we still observed no activation of Rac1 upon treatment with SDF1α (Fig. 3f and Supplementary Fig. 3a-c). This outcome is consistent with PTEN (C124S) inhibiting P-Rex2 by occluding Gβγ binding. In contrast, the EGF-stimulated increase in Rac1 activity was restored in HEK293 cells co-transfected with the catalytically dead PTEN (C124S), compared to the wild-type PTEN (Fig. 3d,f). These data indicate that RTK-activation of P-Rex2 can be inhibited by PTEN increasing the hydrolysis of PI(3,4,5)P_3_.

### The PTEN PDZ-BM and the P-Rex2 PDZ2 domain coordinate PTEN:P-Rex2 complex formation

Despite significant sequence and functional similarity between P-Rex homologues, PTEN only binds and inhibits P-Rex2 but not P-Rex1^3^. As expected, we confirmed that PTEN does not bind P-Rex1 by SPR (Fig. 4a, Supplementary Table 2). This behaviour provided us with an opportunity to delineate further the P-Rex2 sequence regions important for PTEN:P-Rex2 co-inhibition. We designed and purified eight ten-residue P-Rex1 to P-Rex2 sequence-swap mutants that targeted non-conserved sequence regions at each of the three identified PTEN:P-Rex2 crosslinking sites in P-Rex2 (Fig. 4b,c).

**Fig. 4.**
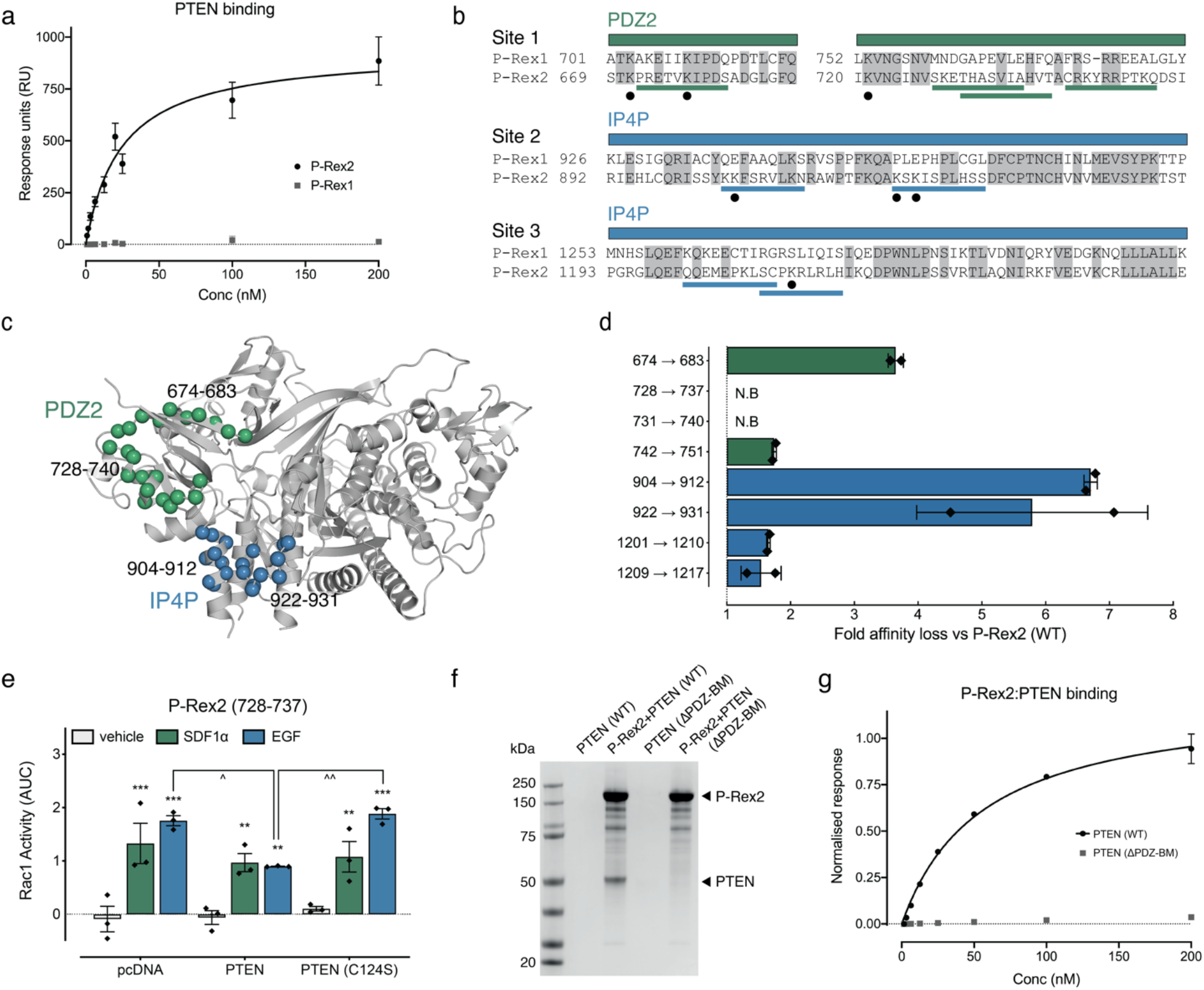
The PTEN PDZ-BM and the P-Rex2 PDZ2 domain are essential for PTEN:P-Rex2 complex formation and Rac1 inhibition. **a**, SPR steady-state response curves of PTEN binding to P-Rex2 or P-Rex1. **b**, Sequence alignments of P-Rex2 PDZ2 and IP4P regions that crosslink to PTEN. Black circles indicate PTEN-crosslinked lysine residues. Solid lines highlight non-conserved regions targeted for sequence-swap mutants. **c**, P-Rex2 DEP2-IP4P domains (modeled from 6PCV^12^) with the location of select sequence-swap mutations highlighted. **d**, Fold decrease of PTEN binding affinity (K_D_) to P-Rex2 sequence-swap mutants compared to P-Rex2 (WT). N.B. indicates no binding. **e**, Rac1 activation upon transient expression of P-Rex2 (728-737) with pcDNA, PTEN, or PTEN (C124S) in HEK293 cells. AUC, area under the curve. Scatter plots show individual data points, symbols/bars represent means, and error bars indicate S.E.M. ** p<0.01 and *** p<0.001 versus vehicle control, ^ p<0.05 and ^^ p<0.01 versus indicated response, two-way ANOVA with Dunnett’s or Tukey’s multiple comparison test. **f**, Coomassie-stained SDS-PAGE of in vitro P-Rex2 pulldowns of recombinant insect-cell purified PTEN (WT) compared to PTEN (ΔPDZ-BM). **g**, SPR steady-state response curves of P-Rex2 binding to PTEN (WT) compared to PTEN (ΔPDZ-BM).

SPR analysis demonstrated that switching residues 674 to 683, within the PDZ2 region, resulted in a four-fold decrease in PTEN:P-Rex2 affinity (Fig. 4d, Supplementary Fig. 4a, Supplementary Table 2). Most notably, we observed a complete loss of binding for two overlapping sequence swap mutants within PDZ2 (728-737 and 731-740), with no significant change in the binding affinity of the final PDZ2 mutant located slightly upstream (742-751) (Fig. 4d, Supplementary Fig. 4a, Supplementary Table 2). Within the IP4P domain, two sequence-swap mutants (residues 904-912 and residues 922-931) reduced the affinity seven- and six-fold, respectively (Fig. 4d, Supplementary Fig. 4a, Supplementary Table 2). The two final IP4P sequence-swap mutants (1201-1210 and 1209-1217) failed to affect the affinity of P-Rex2 for PTEN compared to the wild-type control, suggesting that although this region is close to PTEN in the PTEN:P-Rex2 complex, it does not directly contribute to binding (Fig. 4d, Supplementary Fig. 4a, Supplementary Table 2). Together, these data further narrow-down the PTEN binding site to two non-sequential PDZ2 (674-683, 728-740) and IP4P (904-912, 922-931) sequence regions that cluster at the same surface of the P-Rex2 model (Fig. 4c).

To further confirm the principal role of the P-Rex2 PDZ2 and IP4P domains in PTEN binding, we purified an N-terminally truncated P-Rex2 variant that lacks the DH-PH and DEP1 domains (DEP2-IP4P; residues 472-1606) and measured binding to PTEN by SPR (Supplementary Fig. 4b). We detected no major difference in PTEN affinity for N-terminally truncated P-Rex2 DEP2-IP4P compared to full-length P-Rex2 (Supplementary Table 2). In addition, we observe no binding for the isolated P-Rex2 DH-PH domains to PTEN (Supplementary Fig. 4c). This result is consistent with there being no requirement of the P-Rex2 DH-PH and DEP1 domains for PTEN:P-Rex2 binding, despite minor crosslinks indicating the domains are near the PTEN:P-Rex2 interface.

To examine the effect of uncoupling PTEN and P-Rex2 on Rac1 activation, we transfected HEK293 cells with P-Rex2 (728-737) which was unable to bind PTEN by SPR (Supplementary Fig. 4d-f). Similar to wild-type P-Rex2, we observed an increase in Rac1 activity following stimulation of the cells with both SDF1α and EGF (Fig. 4e and Supplementary Fig. 4g). There was no effect of PTEN co-transfection on the increase in Rac1 activity induced downstream of GPCR stimulation, however there was a small decrease in Rac1 activity induced downstream of RTK stimulation (Fig. 4e and Supplementary Fig. 4h). Given that P-Rex2 (728-737) cannot directly bind PTEN (Fig. 4d), this is likely due to PTEN increasing the hydrolysis of PI(3,4,5)P_3_. This is further supported by the significant increase in Rac1 activity in response to EGF following co-transfection of catalytically dead PTEN (C124S) compared to wild-type PTEN (Fig. 4e and Supplementary Fig. 4h,i). Taken together, these data suggest a dual mode of inhibition of wild-type P-Rex2 by PTEN: whereby PTEN both increases PI(3,4,5)P_3_ hydrolysis, and can directly inhibit the activity of EGF-stimulated P-Rex2 by forming the PTEN:P-Rex2 complex (compare Fig. 3e and Fig. 4e).

We next sought to identify the sequence regions in PTEN that are important for binding P-Rex2. The PTEN tail contains a class-1 PDZ-BM at the extreme C-terminus and would be a logical anchor point for interactions with the P-Rex2 PDZ2 domain. While we observed no crosslinking between the PTEN-tail region and P-Rex2 (Fig 2e), this region contains only a single lysine residue, which decreases the likelihood of detecting a PTEN-tail and P-Rex2 interaction using CLMS. Indeed, previous cellular studies have implicated the PTEN C-terminal tail region in P-Rex2 binding and co-inhibition^4^. We, therefore, sought to investigate the role of the PTEN C-terminus in P-Rex2 binding. We purified a PTEN variant lacking the final four PTEN residues (ΔPDZ-BM) and observed a complete loss of P-Rex2 binding by both SPR and pull-down assays (Fig. 4f,g). Examination of the P-Rex2 PDZ2 domain sequence revealed several residues required for binding class I PDZ-BMs that are not conserved in P-Rex1 (Fig. 5a,b). Residues include a critical histidine (His732) within the PDZ2-binding pocket (which was switched in the P-Rex2 (728-737) and P-Rex2 (731-740) sequence-swap mutants), as well as obligate glycine residues (Gly686, Gly688) of the PDZ carboxylate-binding loop (Fig. 5a,b).

**Fig. 5.**
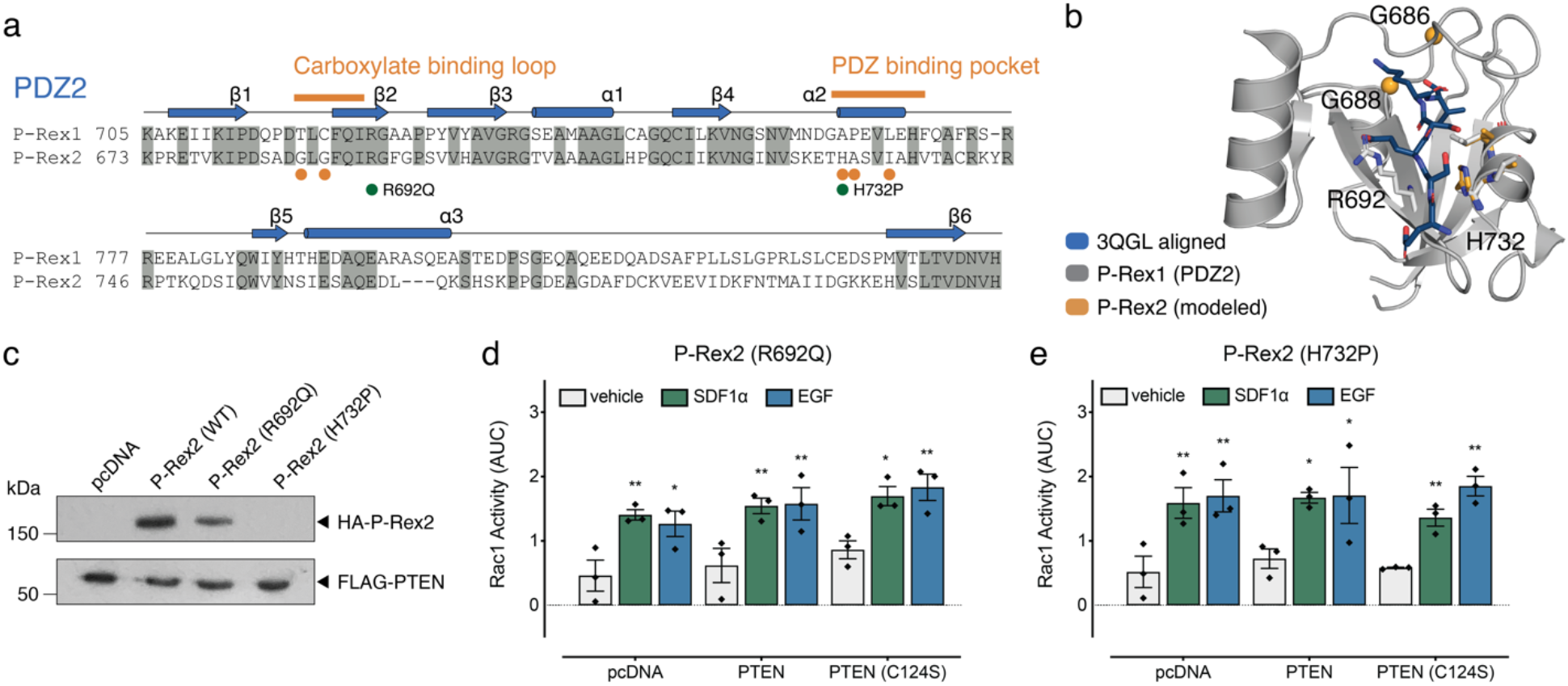
Cancer-associated somatic mutations in the P-Rex2 PDZ2 domain release the PTEN inhibitory effect on Rac1 signaling. **a**, Sequence alignment of the P-Rex1 and P-Rex2 PDZ2 domains. Orange bars indicate PDZ-BM binding pocket residues. Orange spheres indicate crucial PDZ-BM binding residues that are not conserved in P-Rex1. Green spheres highlight the location of the analysed PDZ2 cancer-associated somatic mutants. **b**, The type-1 PDZ-BM from PDB 3QGL^40^ modeled to the binding pocket of the P-Rex1 PDZ2 domain. Residues crucial for binding PDZ-BMs that are conserved in P-Rex2, but not P-Rex1, are highlighted in orange. The location of cancer-associated mutations within the PDZ2 domain are indicated. **c**, FLAG-PTEN pull-down of P-Rex2 (WT) and indicated cancer-associated P-Rex2 mutants. **d-e**, Rac1 activity upon transient expression of indicated P-Rex2 somatic mutants with pcDNA, PTEN, or PTEN (C124S) in HEK293 cells. AUC, area under the curve. Scatter plots show individual data points, symbols/bars represent means, and error bars indicate S.E.M. * p<0.05 and ** p<0.01 versus vehicle control, two-way ANOVA with Dunnett’s multiple comparison test.

Together, these data build a molecular rationale for the selective binding of PTEN to P-Rex2 over P-Rex1, whereby the PTEN C-terminal PDZ-BM binds to the conserved P-Rex2 PDZ2 binding pocket. This anchors PTEN across the P-Rex2 PDZ2 and IP4P domains and inhibits the activation of P-Rex2.

### Cancer-associated P-Rex2 somatic mutations uncouple PTEN inhibition of P-Rex2 to drive unregulated Rac1 activity

Given the critical role of the P-Rex2 PDZ2 domain in PTEN:P-Rex2 binding and co-inhibition, we rationalised that cancer-associated somatic mutations in the P-Rex2 PDZ2 domain may drive Rac1 activation by preventing the assembly of the co-inhibitory PTEN:P-Rex2 complex. We first investigated two somatic mutants with non-synonymous mutations in the PDZ2 domain: P-Rex2 (R692Q), with a mutation located at strand-β2 of the carboxylate binding loop, has been found in skin, endometrial, and breast cancers (COSV55777314^27^), and P-Rex2 (H732P), with a mutation of the critical histidine of the PDZ binding pocket (H732P), which has been reported in breast cancer (COSV55757958^27^) (Fig. 5a,b). FLAG-tag PTEN pulldown experiments in Expi293 cells (an in-suspension variant of HEK293 cells) determined that P-Rex2 (R692Q) mutation reduced PTEN binding by ~60% with the non-conservative P-Rex2 (H732P) mutation eliminating PTEN binding (Fig. 5c). Overexpression of the P-Rex2 (R692Q) and P-Rex2 (H732P) mutants in HEK293 cells resulted in an increase in Rac1 activity in response to both SDF1α and EGF, comparable to wild-type P-Rex2 (Fig. 5d,e and Supplementary Fig. 5). However, unlike wild-type P-Rex2, co-transfection of either PTEN or PTEN (C124S) failed to inhibit Rac1 activation downstream of either SDF1α or EGF stimulation (Fig. 5d,e and Supplementary Fig. 5). These data are consistent with somatic mutations in the PDZ2 domain, allowing P-Rex2 to bypass inhibition by PTEN within the cell. This behaviour, therefore, facilitates unregulated Rac1 activity downstream of an RTK or GPCR stimulus.

Many of the P-Rex2 cancer-associated somatic mutants studied to date are nonsense mutations resulting in the production of truncated P-Rex2 protein products^13,15^. Of these, we selected P-Rex2 (E824*) and P-Rex2 (Q1430*) melanoma-associated truncation mutations for further investigation. Each mutant retains the critical PDZ2 domain required for PTEN binding. However, given the extent of the IP4P domain deletions, it is unlikely that either mutant retains the IP4P:PTEN or the Gβγ binding sites (Fig. 6a,b). Our CLMS data also suggest that neither mutant is likely to be autoinhibited via P-Rex2 intramolecular interactions in the basal state (e.g. Fig. 1d).

**Fig. 6.**
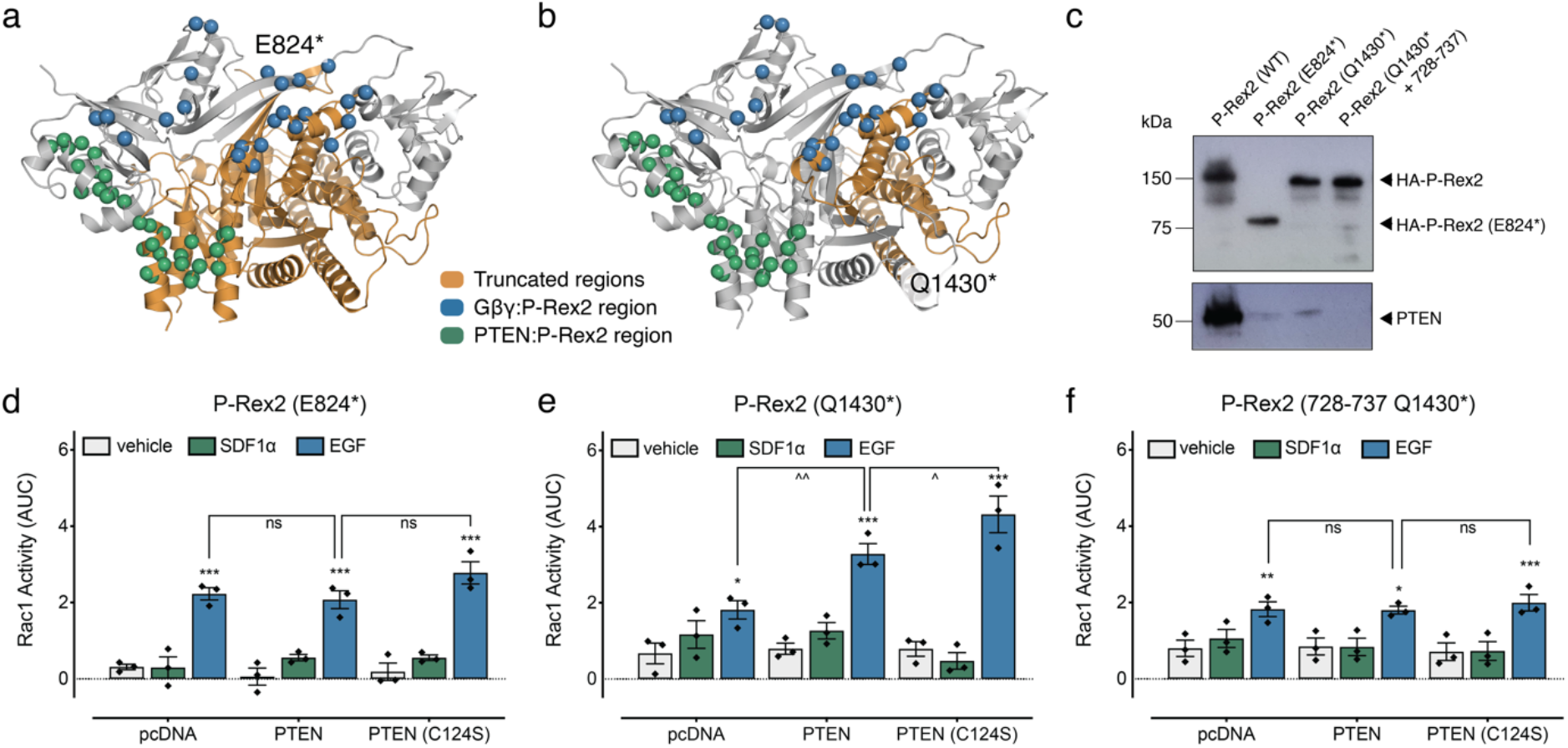
PTEN and P-Rex2 somatic cancer cancer-associated mutations synergize to drive Rac1 activation. **a-b**, P-Rex2 DEP2-IP4P domains (modeled from 6PCV^12^) highlighting the location of truncation mutations to the Gβγ and PTEN interaction sites. **c**, HA-P-Rex2 pulldown of PTEN with P-Rex2 mutants indicated. **d-f**, Rac1 activity upon transient expression of indicated P-Rex2 mutants with pcDNA, PTEN, or PTEN (C124S) in HEK293 cells. Scatter plots show individual data points, symbols/bars represent means, and error bars indicate S.E.M. * p<0.05, ** p<0.01 and ***p<0.001 versus vehicle control; ns, not significant, ^ p<0.05 and ^^ p<0.01 versus indicated response, two-way ANOVA with Dunnett’s or Tukey’s multiple comparison test.

HA-tag P-Rex2 pulldowns in Expi293 cells demonstrated that both P-Rex2 (E824*) and P-Rex2 (Q1430*) had a heavily reduced yet still significant capacity to bind PTEN (Fig. 6c). This binding is consistent with the retention of the PDZ2:PTEN binding site and loss of the IP4P:PTEN binding site. In HEK293 cells transfected with either P-Rex2 (E824*), or P-Rex2 (Q1430*), we also observed a marked reduction in the expression levels of the mutants (Supplementary Fig. 6a). Importantly, expression levels did not change upon PTEN, or PTEN (C124S) co-expression, allowing us to compare Rac1 activity within each P-Rex2 mutant upon PTEN co-expression (Supplementary Fig. 6a-c). Overexpression of P-Rex2 (E824*) resulted in an increase in Rac1 activity following EGF stimulation (Fig. 6d and Supplementary Fig. 6d-f). However, there was a complete loss of SDF1α-stimulated Rac1 activity (Fig. 6d and Supplementary Fig. 6d-f). These data are compatible with the loss of the Gβγ binding site upon IP4P truncation (Fig. 6a). Despite retaining the PDZ2 domain, co-transfection of PTEN, or PTEN (C124S) failed to inhibit the EGF response (Fig. 6d and Supplementary Fig. 6d-f). These data are consistent with PTEN retaining some binding P-Rex2 but failing to inhibit its GEF activity. We suggest that PTEN fails to inhibit P-Rex2 (E824*) because it can no longer stabilise an autoinhibitory basal P-Rex2 fold due to truncation of the IP4P domain.

In HEK293 cells expressing the P-Rex2 (Q1430*) truncation mutant, there was no change in Rac1 activity following stimulation with SDF1α, again consistent with truncation of the Gβγ binding site (Fig. 6e and Supplementary Fig. 6g-i). Rac1 activity was still increased following stimulation with EGF, comparable to wild-type P-Rex2 and P-Rex2 (E824*) (Fig. 6e and Supplementary Fig. 6g-i). Strikingly, however, co-transfection of PTEN, or PTEN (C124S), caused an additional increase in Rac1 activity following stimulation of the cells with EGF (Fig. 6e and Supplementary Fig. 6g-i).

We next sought to test the requirement of PTEN:P-Rex2 complex formation for the enhanced Rac1 response by P-Rex2 (Q1430*). We transfected HEK293 cells with a P-Rex2 (728-737 + Q1430*) variant that combined truncation and PTEN-binding mutations and measured Rac1 activation (Fig. 6f and Supplementary Fig. 6j-l). As expected, we retained the EGF-induced increase in Rac1 activity. However, there is no further increase in Rac1 activation upon co-transfection with PTEN or PTEN (C124S) (Fig. 6f and Supplementary Fig. 6j-l). As expected, HA-P-Rex2 pulldowns in Expi293 cells demonstrated no binding between P-Rex2 (Q1430* + 728-737) and PTEN (Fig. 6c). This behaviour indicates that PTEN binding to P-Rex2 (Q1430*) is critical for the observed increase in Rac1 activity in response to EGF. We suggest that removing the P-Rex2 IP4P regions may free the PTEN C2-membrane binding loops allowing PTEN to localise P-Rex2 (Q1430*) at the membrane. However, other mechanisms, including PTEN stabilisation of truncated P-Rex2, could also play a role.

Together, these data illustrate how somatic P-Rex2 mutations can flip the role of the PTEN:P-Rex2 complex from inhibitory to stimulatory, hyperactivating downstream Rac1 signaling, and potentially contributing to cancer progression.

## DISCUSSION

The co-inhibitory PTEN:P-Rex2 complex forms a central signaling node linking the activation of GPCRs and RTKs to the regulation of both PI3K-AKT and RhoGTPase signaling pathways. Our work provides mechanistic insight into the assembly of the PTEN:P-Rex2 complex as well as providing a rationale for the dysregulation of this central signaling node in cancer. We propose a two-stage binding model for PTEN:P-Rex2 complex assembly (Fig. 7a). Binding is first initiated by an interaction between the PTEN PDZ-BM tail and the P-Rex2 PDZ2 binding pocket and then stabilised by extensive interactions between residues in the PTEN C2-DUSP domains and two key binding sites in P-Rex2 covering a continuous surface formed by the PDZ2 and IP4P domains.

**Fig. 7.**
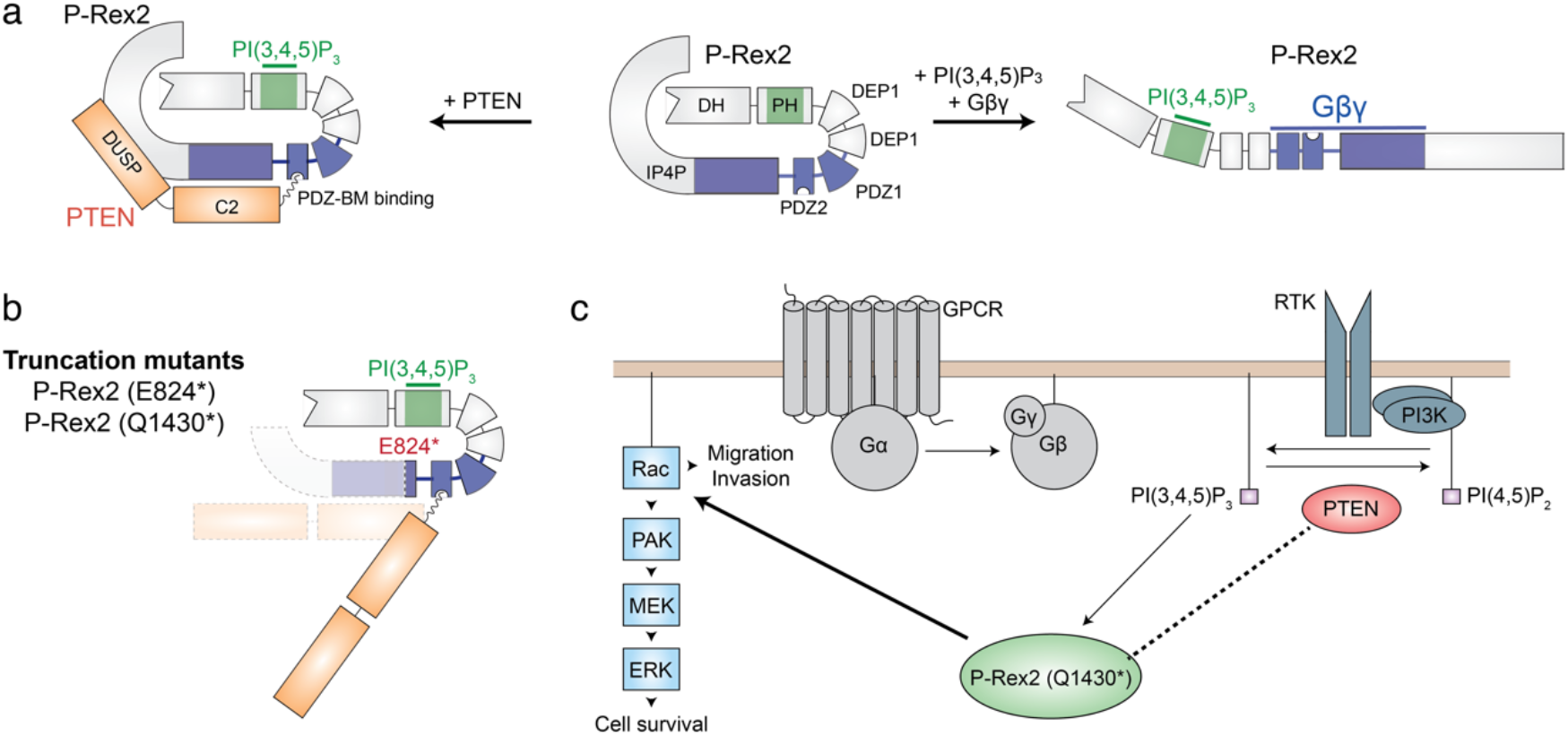
PTEN:P-Rex2 complex assembly and dysregulation by cancer-associated mutations. **a**, Schematic illustration of assembly and co-inhibition of the PTEN:P-Rex2 complex. The basal autoinhibited P-Rex2 conformation is stabilised by interactions between the DH-PH and IP4P domains (centre). PTEN:P-Rex2 assembly is initiated by binding between the PTEN PDZ-BM tail and the P-Rex2 PDZ2 binding pocket and then stabilised by extensive interactions between residues in the PTEN C2-DUSP domains and two critical binding sites in the P-Rex2 PDZ2 and IP4P domains (left). This interaction locks P-Rex2 in the autoinhibited state preventing activation by Gβγ (right). **b**, P-Rex2 truncation mutations remove the IP4P:DH-PH interface and the Gβγ-binding site. **c**, Loss of the Gβγ-binding site prevents P-Rex2 activation downstream of GPCR stimulus. However, truncated P-Rex2 is still activated downstream of RTK stimulation by an increase in PI(3,4,5)P_3_ levels. Truncation mutations retain low-affinity PTEN binding (dotted line), but GEF activity is no longer inhibited by PTEN:P-Rex2 complex assembly. Strikingly, co-transfection of PTEN, or PTEN (C124S), causes an additional increase in Rac1 activity downstream of RTK stimulus.

We find that the anchoring of the PTEN PDZ-BM tail within the P-Rex2 PDZ2 binding pocket likely initiates the formation of the PTEN:P-Rex2 complex. Multiple lines of evidence converge to support this model. PDZ2 residues required to bind class I PDZ-BMs are conserved in P-Rex2 compared to P-Rex1 (which cannot bind PTEN), and PTEN:P-Rex2 binding is lost in PDZ2 P-Rex1-sequence-swap mutants. Further, cancer-associated somatic mutations within the P-Rex2 PDZ2 domain prevent PTEN from inhibiting Rac1 signaling downstream of GPCR and RTK activation. Finally, truncation of the PTEN PDZ-BM prevents P-Rex2 binding in vitro. Interestingly, mutations that result in deletion of the PTEN PDZ-BM have been identified in tumours^4^, and our model predicts that these mutations may promote tumour growth by preventing PTEN:P-Rex2 binding, therefore preventing PTEN inhibition of P-Rex2. This first stage of our model is also in agreement with the majority of coIP and pulldown studies that demonstrate necessity of the PTEN PDZ-BM for binding to P-Rex2^3,4,13,14^ and the requirement of an intact PTEN PDZ-BM for PTEN-mediated suppression of P-Rex2-dependent cell invasion^4^.

PDZ:PDZ-BM interfaces are usually low affinity^28^, and this is compatible with an additional role for the P-Rex2 PDZ2-IP4P surface region in promoting complex assembly. Using CLMS, we identified extensive crosslinks between P-Rex2 and the PTEN C2 domain. Critically, PTEN binding to P-Rex2 was reduced in P-Rex2 mutants where sections of the IP4P domain sequence were swapped with the corresponding region from P-Rex1. Together, this suggests that the PTEN C2 domain and possibly the PTEN DUSP domain form a stabilising bridge across the P-Rex2 PDZ2-IP4P surface. Given the importance of the P-Rex2 PDZ2-IP4P surface region for PTEN binding, it is noteworthy that isolated P-Rex2 PDZ2-binding pocket mutations (both synthetic and somatic) can completely inhibit PTEN binding. These data are consistent with our two-stage binding model, whereby anchoring of the PTEN PDZ-BM tail region to the P-Rex2 PDZ2-binding pocket exposes, or positions, PTEN DUSP-C2 residues for binding to the P-Rex2 surface. Previous studies suggest the P-Rex2 DH-PH domains directly interact with PTEN to inhibit its phosphatase activity^3,14^. However, we were not able to detect an extensive interaction between these regions by CLMS, or any binding between the isolated P-Rex2 DH-PH domains and PTEN by SPR. Instead, we found that the P-Rex2 DEP2-IP4P domains alone were sufficient to retain the full PTEN binding affinity. Nevertheless, an interface may exist between PTEN and the P-Rex2 DH-PH domains in a cellular context, especially given the predisposition of both PTEN and P-Rex2 to form multiple conformations^5,29^.

The use of CLMS, and measurements of Rac1 activity in live cells, also allowed us to delineate the mechanism underlying the dual inhibition of PTEN and P-Rex2 in the PTEN:P-Rex2 complex. Rather than PTEN binding directly to the P-Rex2 DH-PH domains to inhibit GEF activity, we propose a dual model of P-Rex2 inhibition. PTEN both occludes Gβγ binding to P-Rex2 by binding across the same surface (and therefore inhibits GPCR-mediated changes in Rac1 activity), and allosterically promotes an autoinhibited P-Rex2 conformation. The autoinhibitory mechanism of P-Rex family proteins has proven difficult to ascertain, beyond a long-known requirement for the multiple C-terminal P-Rex domains^10,29^. Here, CLMS provides direct evidence for a role of the IP4P domain, including the loop region, in occluding Rac1 binding to the DH-PH domain, thereby providing a mechanism for P-Rex2 autoinhibition. Interestingly, PAK kinases are known to inhibit membrane-bound P-Rex2 by phosphorylating this IP4P loop region (e.g. Ser1107) downstream of IGF signaling^30^. Phosphorylation of the exposed IP4P loop within membrane-bound active P-Rex2 may promote the autoinhibitory DH-PH:IP4P domain interactions identified in our model, resulting in the inhibition of P-Rex2 GEF activity.

The mechanism whereby P-Rex2 inhibits PTEN appears more straightforward. Using CLMS we identify the PTEN Cα2 helix and C2 membrane-binding loops, which are critical for association of PTEN with the membrane, coordinated toward the PTEN:P-Rex2 interface. This is in agreement with PTEN truncation studies which have highlighted a role for the PTEN C2 domain in binding to P-Rex2, in addition to the PTEN PDZ-BM tail^4^. Together with liposome pulldown experiments, these data indicate that assembly of the PTEN:P-Rex2 complex causes inhibition of PTEN by preventing both PTEN PI(3,4,5)P_3_ hydrolysis and membrane association.

Using our models of PTEN:P-Rex2 binding and co-inhibition, we were able to characterise a new class of gain-of-function cancer-associated P-Rex2 mutations. These mutations occur in the P-Rex2 PDZ2 domain, which allows P-Rex2 to retain GEF activity downstream of both GPCR and RTK stimuli yet avoid regulation by PTEN inhibition. This class of mutation ultimately results in an unregulated increase in cellular P-Rex2 activity, and therefore RhoGTPase activity (e.g. Rac1). Other somatic mutations within the PTEN:P-Rex2 interface, or mutations that misfold or truncate the PDZ2 domain, would likely also uncouple PTEN regulation from RhoGTPase signaling and belong to the same gain-of-function mutant class. Although not shown here, in addition to unregulated activation of RhoGTPase signaling, loss of PTEN:P-Rex2 binding would likely release PTEN to inhibit PI3K-AKT signaling instead.

The second class of cancer-associated P-Rex2 mutations are no longer activated downstream of GPCRs but can be hyperactivated by RTKs (Fig. 7b,c). These mutants have large truncations within the P-Rex2 IP4P domain, which deletes the Gβγ binding site^12^ but retains the PI(3,4,5)P_3_-binding PH domain that coordinates EGF-induced activity and membrane association^9^. Our CLMS data suggest that truncation, or mutation-induced misfolding, of significant regions of the IP4P domain would also hyperactivate P-Rex2 by disrupting the intramolecular associations that are required for autoinhibition. This is consistent with increased Rac1 activation by truncated P-Rex2 in immortalised melanocytes or xenograft tumour-derived cells^13^. We also demonstrate truncated P-Rex2 mutants evade PTEN inhibition downstream of EGF stimulation, providing an additional explanation for the observed increase in Rac1 activation upon P-Rex2 truncation.

Interestingly, although we observe a loss of PTEN inhibition for this second class of mutants, coimmunoprecipitation and pull-down experiments demonstrate that truncated P-Rex2 retains PTEN binding. This binding likely represents the first stage of the PTEN:P-Rex2 interaction, mediated by the intact P-Rex2 PDZ2 domain. Interestingly, previous studies have shown that ectopically expressed truncated P-Rex2 is unable to bind endogenous PTEN^13^. Given the loss of the extensive P-Rex2 PDZ2-IP4P:PTEN binding surface upon P-Rex2 truncation, we would expect a decrease in PTEN:P-Rex2 binding affinity with binding reliant only on the initial P-Rex2 PDZ2-binding pocket:PTEN PDZ-BM interaction. As such, we can observe this low-affinity interaction in experiments with high levels of PTEN (e.g. coimmunoprecipitation with transfected proteins), but not in systems with low PTEN levels (e.g. coimmunoprecipitation with endogenous PTEN^13^).

Most strikingly, Rac1 activation by P-Rex2 (Q1430*) is enhanced upon expression of either PTEN or PTEN (C124S). This increase is well above the level of Rac1 activation observed without PTEN expression and appears reliant on the PDZ2:PDZ-BM interaction. We suggest PTEN binds P-Rex2 (Q1430*) but can no longer allosterically support a closed, autoinhibited P-Rex2 conformation without a folded P-Rex2 IP4P domain. PTEN binding may, however, increase the stability of truncated P-Rex2 (and therefore its Rho GEF activity) or could enhance membrane localisation of the PTEN:P-Rex2 complex, perhaps allowing more efficient activation of RhoGTPase signaling pathways. This hyperactivity is particularly interesting given that P-Rex2 somatic mutations are generally associated with high PTEN expression^4^. As such, the coincidence of P-Rex2 (Q1430*) with high PTEN expression, or catalytically-dead PTEN cancer-associated mutations (e.g. C124S and G129E), in tumours, may drive Rac1 activation further than complete PTEN loss. Whether P-Rex2 and PTEN cancer-associated mutations synergise to drive metastasis require further investigation.

While a complete understanding of the intricate functional interdependency between the PTEN tumour suppressor and the P-Rex2 oncogene will require a high-resolution structure of the PTEN:P-Rex2 complex, this remains a significant challenge given the conformation flexibility of both full-length PTEN and P-Rex2. Here we use CLMS, mutagenesis, and single-cell Rac1 measurements to provide a much-needed mechanistic framework to understand the frequent dysregulation of the PTEN:P-Rex2 complex that occurs in metastatic cancer. We propose a two-stage model of PTEN:P-Rex2 binding, initiated by the PTEN PDZ-BM tail and the P-Rex2 PDZ2 binding pocket, and then stabilised by an extensive interface provided by the PTEN DUSP-C2 domains and a large surface on P-Rex2 formed by the IP4P and PDZ2 domains. The PTEN:P-Rex2 complex inhibits P-Rex2 by both occluding Gβγ binding and allosterically promoting P-Rex2 autoinhibition. The PTEN:P-Rex2 interface also occludes the PTEN PI(3,4,5)P_3_-binding pocket and C2 domain membrane-binding loops, therefore inhibiting PTEN. More broadly, this study highlights the application of CLMS as a powerful tool capable of providing much-needed mechanistic insights that extend beyond large protein complexes to small, or highly mobile interfaces.

## METHODS

### Cloning

N-terminally His-tagged P-Rex2 was synthesised and cloned into pFastBac1 (Invitrogen). His-tagged PTEN, or PTEN (ΔPDZ-BM) were synthesised and cloned into pFastBac1 (Invitrogen). For Gβγ expression, Gβ_1_ and His-Gγ_2_ (C68S) were synthesised and cloned into pFastBac Dual (Invitrogen). For mammalian expression, N-terminally HA-tagged P-Rex2, or N-terminally Flag-tagged PTEN, were synthesised and cloned into pcDNA3.1 (Invitrogen). All P-Rex2 and PTEN mutations were synthesised and sequence-verified in the same vectors by Genscript. Rac1 (G12V) (residues 1–177) was cloned into pGEXTEV, a modified version of pGEX-4T-1 (GE Healthcare) where the thrombin site is replaced with a TEV cleavage site.

### Protein expression and purification

His-tagged P-Rex2 was expressed in Sf9 cells for 2 days following baculovirus infection (Bac-to-Bac, Invitrogen). Cells were harvested by centrifugation and lysed by sonication in 20 mM Tris-HCl pH 8.0, 500 mM NaCl, 10% (v/v) glycerol, and 5 mM β-mercaptoethanol. Lysates were cleared by centrifugation, filtered at 0.45 μm, and incubated with Ni-NTA resin (Qiagen) and 20 mM imidazole at 4 °C with agitation. The resin was washed with lysis buffer, and P-Rex2 was eluted with 500 mM imidazole in the same buffer. Protein was dialysed overnight into 20 mM Tris-HCl pH 8.0, 200 mM NaCl, 10% (v/v) glycerol, 2 mM DTT, and 5 mM MnCl_2_. λ-phosphatase was added to the protein before dialysis. Protein was diluted 1:10 in 20 mM Tris-HCl pH 8.0, 50 mM NaCl, 2 mM DTT, and 5% (v/v) glycerol for further purification by anion exchange on a Mono Q 5/50 GL column (GE Healthcare) with a gradient from 50 mM to 1 M NaCl. P-Rex2-containing fractions were further purified on a Superdex 200 16/60 size-exclusion column (GE Healthcare) equilibrated in 20 mM HEPES pH 8.0, 150 mM NaCl, and 2 mM DTT. All P-Rex2 variants, PTEN (WT) and PTEN (ΔPDZ-BM) were expressed and purified as per P-Rex2 (WT). The PTEN:P-Rex2 complex was purified by incubating a 3:1 molar ratio of PTEN to P-Rex2 in 20 mM HEPES pH 8.0, 150 mM NaCl, and 2 mM DTT for 1 hour prior to purification on a Superdex 200 10/300 size-exclusion column (GE Healthcare). Rac1 (G12V) was purified as previously described^10^. His-tagged Gβγ was expressed in Sf9 cells for 2 days following baculovirus infection (pFastBacDual, Bac-to-Bac, Invitrogen). Cells were harvested by centrifugation and lysed by sonication in 20 mM Tris-HCl pH 8.0, 500 mM NaCl, 1 mM MgCl_2_, and 5 mM β-mercaptoethanol. Lysates were cleared by centrifugation, filtered at 0.45 μm, and incubated with Ni-NTA resin (Qiagen) and 20 mM imidazole at 4 °C with agitation. The resin was washed with lysis buffer, and Gβγ was eluted with 300 mM imidazole in the same buffer. Protein was diluted 1:10 in 20 mM Tris-HCl pH 8.0, 80 mM NaCl, and 2 mM DTT for further purification by anion exchange on a Mono Q 5/50 GL column (GE Healthcare) with a gradient from 80 mM to 1 M NaCl. Gβγ was further purified on a Superdex 75 16/60 size-exclusion column (GE Healthcare) in 20 mM HEPES pH 8.0, 150 mM NaCl, and 2 mM DTT.

### Phosphatase-activity assay

Samples were incubated in 20mM Tris-HCl pH 8.0, 150 mM NaCl, and 2mM DTT at 37 °C for 30 min prior to the addition of 15 μM di-C8-D-myo-phosphatidylinositol 3,4,5-triphosphate (Echelon). After which, reactions were incubated at 37 °C for 5 min and stopped by the addition of malachite green reagent (Sigma Aldrich). Absorbance was measured at 620 nm on a FLUOstar Omega plate reader (BMG).

### ^35^S-GTPγS binding assay

Rac1 was loaded with GDP by incubating 12.5 μM His-Rac1 with 20 μM GDP and 5 mM EDTA in 20 mM HEPES pH 7.4, 100 mM NaCl, and 1 mM ethylene glycol-bis(β-aminoethyl ether)-N,N,N’,N’-tetraacetic acid for 20 min at room temperature. 20 mM MgCl_2_ was then added to facilitate nucleotide loading. In parallel, 300 nM Gβγ and liposomes (liposome composition as per^31^) were equilibrated at room temperature for 20 min in 20 mM HEPES pH 7.4, 100 mM NaCl, 1 mM EGTA, 5 mM MgCl_2_, 5 mM DTT, and 50 μg/ml bovine serum albumin. Subsequently, 100 nM GDP-loaded His-Rac1 was added to the Gβγ/liposome samples and incubated for a further 10 min at room temperature. 50 nM P-Rex2 and 5 μM ^35^S-GTPγS were added to the reaction and incubated at room temperature for a further 15 min. Reactions were stopped by the addition of 20 mM HEPES pH 7.4, 100 mM NaCl, 10 mM MgCl_2_, 10 mM imidazole and Ni-NTA resin prior to incubation at 4 °C on a rotary mixer. Resin was washed three times prior to the addition of Ultima Gold scintillation cocktail (PerkinElmer). Radioactivity was measured with a Tri-Carb 2900TR liquid scintillation counter (PerkinElmer).

### Analytical size-exclusion chromatography

To demonstrate PTEN:P-Rex2 complex formation, a 3:1 molar ratio of PTEN (30 μM) to P-Rex2 (WT) (10 μM) was incubated at room temperature for 1 hour in 10 mM HEPES, 150 mM NaCl, and 2 mM DTT prior to injection onto a Superdex 200 10/300 GL column (GE Healthcare). PTEN and P-Rex2 alone at corresponding concentrations demonstrate the peak-shift upon complex formation.

### CLMS

5 μM PTEN, P-Rex2, or the PTEN:P-Rex2 complex in 20 mM HEPES pH 8.0, 150 mM NaCl, and 2 mM DTT were incubated at a 1:100 molar ratio with either BS^3^ (Thermo Fisher) or BS^2^G (Thermo Fisher) crosslinkers. For BS^3^, samples were incubated at room temperature for 20 min prior to the addition of 50 mM Tris-HCl pH 8.0 to quench the reaction. For BS^2^G, samples were incubated at room temperature for 30 min prior to the addition of 20 mM NH_4_HCO_3_ to quench the reaction. For both crosslinkers, samples were snap-frozen in liquid nitrogen prior to further processing. Samples were then denatured for 30 min at 65°C in the presence of 10 mM DTT. 40 mM chloroacetamide was added to the samples prior to incubation for 20 min at room temperature in the dark. A 1:100 (w:w) ratio of trypsin was added to the samples and further incubated at 37°C overnight. Digestion was stopped upon the addition of 1% (v/v) formic acid. Samples were cleaned using OMIX C18 pipette tips (Agilent Technologies) and stored in 0.1% formic acid prior to mass spectrometry. The samples were analysed by LC-MS/MS using a Dionex Ultimate 3000 RSLCnano system coupled onto an Orbitrap Fusion Tribrid instrument (Thermo Fisher). An Acclaim PepMap RSLC analytical column (75 μm × 50 cm, nanoViper, C18, 2 μm, 100Å; Thermo Scientific) and an Acclaim PepMap 100 trap column (100 μm × 2 cm, nanoViper, C18, 5 μm, 100Å; Thermo Scientific) were used to separate tryptic peptides by increasing concentrations of 80% acetonitrile (can) / 0.1% formic acid at a flow of 250 nl/min for 90 min. The mass spectrometer was operated in data-dependent mode with the following parameters. The cycle time was controlled for 3 seconds. The MS1 resolution was set at 120,000 and scan range of 375-2000 m/z. The AGC target was set at 1.0e^6^ with an injection time of 118 ms. The MS2 resolution was set at 60,000 and the AGC target was set at 4.0e^5^ with an injection time of 118 ms. pLink and pLink2 were used to identify BS^3^- or BS^2^G-crosslinked peptides^32^. Intramolecular crosslinked peptides were analysed if they had been identified at least twice with a P value of less than 10^−4^. Intermolecular crosslinked peptides were used for analysis if they had been identified once with a P value of less than 10^−4^. Visual representations of crosslinked peptides were generated in Circos^33^. All structural models were generated in SWISS-MODEL^34^ with PDB templates indicated in the main text.

### SPR

SPR experiments were performed on a Biacore T200 (GE Healthcare) in running buffer containing 10 mM HEPES pH 8, 150 mM NaCl, 3 mM EDTA, 0.5 mM DTT, and 0.025% NP-40 at 25 °C with a flow rate of 30 μl/min. To measure binding of PTEN to P-Rex2 (WT), P-Rex1 (WT), and P-Rex2 variants (including sequence-swap and P-Rex2 472-1606), we immobilised PTEN on flow-cell (FC) 2 of a CM4 sensor chip (GE Healthcare) using amine coupling (NHS/EDC). Indicated serial dilutions of the analyte in running buffer were injected for 120 sec, with dissociation monitored for 600 sec, and the sensor surface regenerated using 10 mM NaOH. To measure binding of P-Rex2 to PTEN (WT), and PTEN (ΔPDZ-BM), we immobilised PTEN (WT) and PTEN (ΔPDZ-BM) to FC2 and FC3 of a CM4 chip (GE Healthcare) respectively, by amine coupling (NHS/EDC). Indicated serial dilutions of the analyte in running buffer were injected for 120 secs, with dissociation monitored for 600 secs, and the sensor surface regenerated using 10 mM NaOH. To measure Gβγ binding to P-Rex2, or the PTEN:P-Rex2 complex, FC2 was immobilised with purified Gβγ by amine coupling (NHS/EDC). Indicated serial dilutions of the analyte in running buffer were injected for 120 secs, with dissociation monitored for 600 secs, and the sensor surface regenerated using 10 mM glycine pH 3.5. To measure Rac1 binding to P-Rex2, or the PTEN:P-Rex2 complex, FC3 was immobilised with purified Rac1 by amine coupling (NHS/EDC). Indicated serial dilutions of the analyte in running buffer were injected for 120 secs, with dissociation monitored for 600 secs, and the sensor surface regenerated using 10 mM NaOH. All sensorgrams were FC reference-subtracted before analysis. In each case, we carried out at least two independent analyses, and kinetic constants are reported +/− standard deviation (Supplementary Table 2).

### PTEN PIP_3_-liposome pulldown assays

λ-phosphatase treated P-Rex2 (578-1606), PTEN, or the P-Rex2 (578-1606):PTEN complex were added PIP_3_-liposomes (lipid composition as per^35^) in 25 mM Tris pH 7.5, 150 mM NaCl, 2 mM DTT, and 5% (v/v) glycerol. All tubes were incubated on an orbital roller for 1 hr at 4 °C, then centrifuged (16,000 *g*, 30 min, 4 °C). The supernatant was removed, and the pellet was washed and transfered to a fresh Eppendorf tube to reduce non-specific binding. The centrifugation wash process was repeated a further two times and the supernatant, pellet and final wash samples were analysed by SDS-PAGE.

### Cell culture and transfection

HEK293 cells were maintained in Dulbecco’s Modified Eagle’s Medium (Invitrogen) with 5% (v/v) FBS. Cells were grown in a humidified atmosphere at 37°C with 5% CO_2_. All assay plates were coated with 5 μg/cm^2^ poly-D-lysine prior to use. For verification of P-Rex2 mutant expression by western blotting, cells were transfected with 1.5 μg/well DNA in suspension using 25 kDa linear polyethyleneimine (PEI) at a ratio of 1:6 DNA:PEI and seeded in 6-well plates in complete growth medium. For FRET experiments, cells were transfected with 45 ng/well RaichuEV-Rac1, 50 ng/well P-Rex2 (WT or mutants) and 50 ng/well pcDNA or PTEN (WT or C124S) in suspension using PEI at a ratio of 1:6 DNA:PEI, and seeded in black optically-clear 96-well plates in complete growth medium. Experiments were performed 48 h following transfection.

### Cell lysate preparation and western analysis

Cells were harvested in ice-cold PBS and pellets were resuspended in ice-cold lysis buffer containing 50 mM Tris-HCl pH7.5, 150 mM NaCl, 1 mM MgCl_2_, 1% v/v NP-40, 1% SDS, 1 mM PMSF and protease mini EDTA-free inhibitor cocktail (Roche). Cells were incubated for 30 min on ice and lysed by passing through a 21-gauge needle 10 times. Lysates were centrifuged (200 g, 4°C, 5 min), and supernatants were recovered. For western blotting, the total protein concentration was quantified using a Bradford detection assay. 50 μg total protein was run on 10% SDS-PAGE gels for HEK293 samples. Protein was transferred to nitrocellulose membranes by electroblotting in CAPS buffer before membrane blocking in 5% (w/v) milk powder in Tris-HCl-Buffered Saline (TBS) for 1 hr. Blots were incubated overnight at 4°C with either anti-HA (Abcam, ab9110), anti-PTEN (Invitrogen, 2F4C), or anti-α-tubulin (Cell Signaling Technology 3873), at 1:1000 in 1% (w/v) milk in TBS. Blots were washed 3 times with TBS-T (0.05% v/v Tween-20 in TBS) before incubation with HRP conjugated antibody (Cell Signaling Technology 7076S; anti-mouse-HRP or Cell Signaling Technology; 7074S; anti-rabbit-HRP) at 1:1000 in 2.5% (w/v) milk in TBS for 2 hr. Blots were then washed 3 times in TBS-T and bands visualized with Amersham ECL chemiluminescence reagent (GE Healthcare). Blots were stripped (62.5 mM Tris-HCl pH 6.5, 2% w/v SDS, 50 mM β-mercaptoethanol for 1 hr at 65°C) and re-probed as described with anti-PTEN and finally anti-α-tubulin.

### FRET imaging of live cells

Changes in Rac1 activity were detected using RaichuEV-Rac1^36,37^, which undergoes a conformational change after GTP displaces guanosine diphosphate (GDP) within a target sequence. RaichuEV-Rac1 was from M. Matsuda and K. Aoki (Kyoto University, Japan) and is contained in the pCAGGS vector from J. Miyazaki (Osaka University, Japan). Fluorescence imaging was performed using a high content PerkinElmer Operetta with an Olympus LUCPlanFLN 20x (NA 0.45) objective as previously described^38^, with some modifications. For emission ratio analysis, cells were excited sequentially (410-430 excitation filter) with emission measured using 520-560 and 460-500 emission filters. Cells were imaged every 1 min, allowing image capture of up to 12 wells per minute. Cells were equilibrated in HBSS (Invitrogen) at 37°C, before baseline emission ratio images were captured for 4 min. Cells were challenged with an EC_80_ concentration of SDF1α (30 nM), EGF (10 ng/mL) or vehicle (0.13% w/v BSA in PBS), and images were captured for 20 min. Cells were then stimulated with the positive control (1 μM isoprenaline, 50 ng/mL EGF, 10 μM AlCl_3_ and 10 mM NaF) for 10 min to generate a maximal increase in Rac1 activity, and positive emission ratio images were captured for 4 min. Data were analyzed as described^38^ using in-house automated macros within the FIJI distribution of ImageJ. Cells with >5% change in F/F_0_ (FRET ratio relative to baseline for each cell) after stimulation with positive controls were selected for analysis. Data are expressed as the average emission ratio relative to the maximal response for each cell (F/F_Max_) in a biological replicate, with the area under the curve calculated using GraphPad Prism. Pseudocolor ratiometric images of the cells were generated as described^39^. Data are expressed as the mean ± SEM, of 3 independent experiments. Statistical significance was determined by one-way or two-way ANOVA with Tukey’s or Dunnett’s multiple comparison test, with p<0.05 deemed significant.

## ACKNOWLEDGEMENTS

We would like to thank Mr. Cameron Nowell (Head, Imaging FACS and Analysis Core, Monash Institute of Pharmaceutical Sciences, Monash University) for modifying the automated analysis scripts for use with the Operetta high content imager. C.M.L. was supported by an Australian Postgraduate Award and a Monash Golden Jubilee Postgraduate Research Award. J.C.W. is a National Health and Medical Research Council of Australia (NHMRC) Senior Principal Research Fellow and Australian Research Council (ARC) Laureate Fellow. M.L.H was supported by a NHMRC RD Wright Fellowship (1061687). This research was supported by NHMRC Project Grants to C.A.M. (APP1104614) and A.M.E (APP1146578, APP1128120). We also acknowledge the support of the office of the Vice-Provost for Research and Research Infrastructure (VPRRI) at Monash University and of Bioplatforms Australia (BPA) as part of the National Collaborative Research Infrastructure Strategy (NCRIS).

## AUTHOR CONTRIBUTIONS

L.D., C.M.L., E.A.M., Y-G.C., A.M.E., carried out *in vitro* and *in vivo* experiments. C.H. and R.B.S. conducted mass-spectrometry experiments and data processing. S.C. and M.L.H. conducted single-cell FRET studies and data analysis. All authors contributed to the interpretation of the results and helped write the manuscript.

