## Supplementary Material for "Structural analysis of the PTEN:P-Rex2 signalling node reveals how cancer-associated mutations coordinate to hyperactivate Rac1"

**Supplementary Table 1. CLMS data**
See Supplementary_table_1.xlsx

**Supplementary Table 2. SPR analysis of P-Rex2 variants binding to PTEN.**

| P-Rex2 (variant) | $K_D$ (nM) +/- SD | Fold affinity loss Vs WT |
|---|---|---|
| Wild-type | 24.4 +/- 0.3 | - |
| 674-683 | 88.9 +/- 4.0 | 3.6 |
| 728-737 | No Binding | - |
| 731-740 | No Binding | - |
| 742-751 | 42.4 +/- 0.3 | 1.7 |
| 904-912 | 163.3 +/- 4.5 | 6.7 |
| 922-931 | 140.7 +/- 42.4 | 5.8 |
| 1201-1210 | 40.3 +/- 0.2 | 1.7 |
| 1209-1217 | 37.4 +/- 7.2 | 1.5 |
| 472-1606 | 51.7 +/- 5.7 | 2.1 |

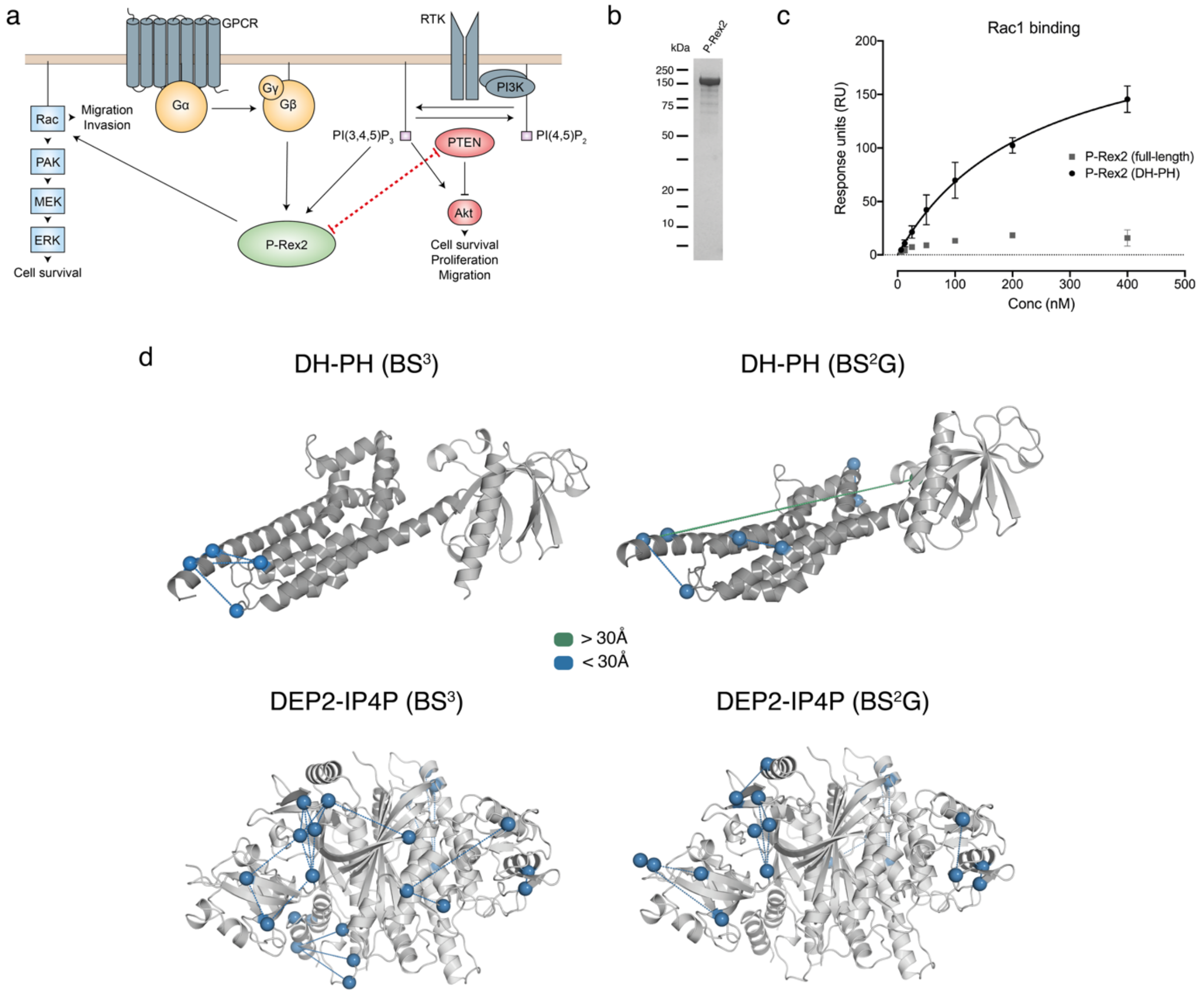


**Fig. S1. Biophysical characterisation of P-Rex2.** **a**, Schematic representation of the PTEN:P-Rex2 signaling pathway. **b**, Coomassie-stained SDS-PAGE of insect-cell purified P-Rex2. **c**, Steady-state response curves of Rac1 binding to full-length P-Rex2 or the isolated P-Rex2 (DH-PH) domains. **d**, CLMS was verified by mapping crosslinks on P-Rex2 models. Blue spheres/lines indicate permissible crosslinks (<30 Å) and green spheres/lines indicate non-permissible (>30 Å).

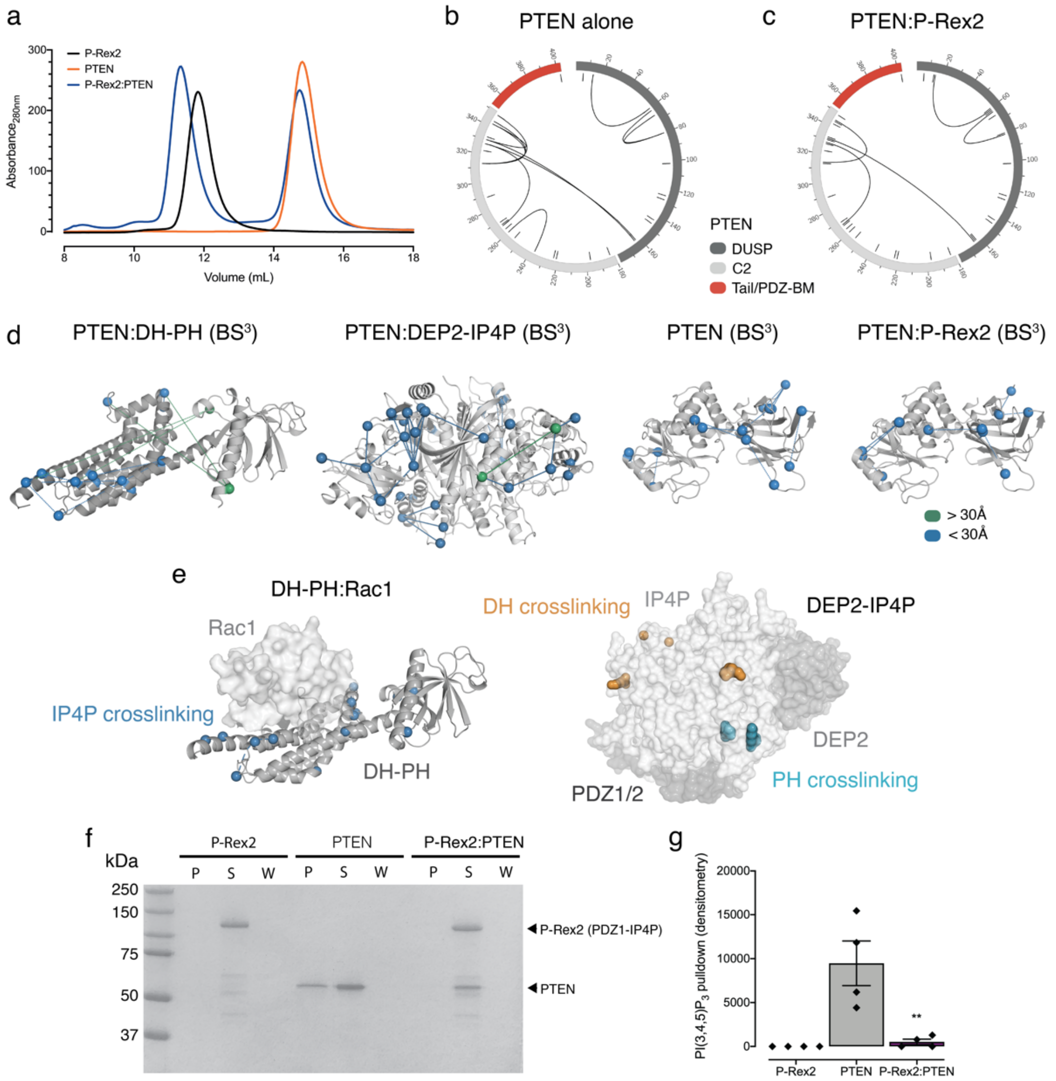


**Fig. S2. Biophysical characterisation of the PTEN:P-Rex2 complex. a**, SEC analysis of P-Rex2, PTEN and the PTEN:P-Rex2 complex. **b-c**, PTEN intramolecular $BS^3$ crosslinks identified by CLMS in the absence (**b**) or presence (**c**) of P-Rex2. Black lines show the identified crosslinks and short black dashes correspond to lysine residues within the primary sequence. **d**, CLMS was verified by mapping crosslinks on P-Rex2 models or the PTEN structure (1D5R[1]). Blue spheres/lines show permissible crosslinks (<30 Å) and green spheres/lines display non-permissible crosslinks (>30 Å). **e**, Intramolecular P-Rex2 crosslinking sites within the PTEN:P-Rex2 complex mapped on the P-Rex2 (DH-PH) domains (modeled from 4YON[2]). Intradomain P-Rex2 crosslinking sites within the PTEN:P-Rex2 complex mapped on the P-Rex2 (DEP2-IP4P) domains (modeled from 6PCV[3]). **f**, PI(3,4,5)$P_3$-liposome pulldown assay of P-Rex2, PTEN, and the PTEN:P-Rex2(PDZ1-IP4P) complex. P equals pellet, S equals supernatant, W equals wash. **g**, Densitometry analysis of the PI(3,4,5)$P_3$-liposome pulldown assay shown in f. ** $p<0.01$ PTEN versus PTEN:P-Rex2, oneway ANOVA with Dunnett's multiple comparison test (n=4).

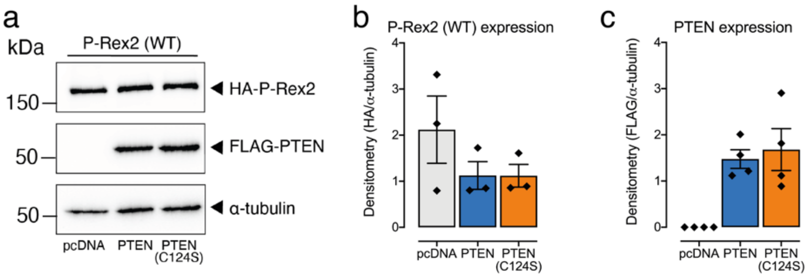


**Fig. S3. Western analysis of P-Rex2 and PTEN expression. a**, Representative western blot showing HA-P-Rex2 and FLAG-PTEN (WT or C124S) expression following transfection in HEK293 cells. **b**, Quantification of HA (P-Rex2) and **c**, FLAG (PTEN) expression levels relative to $\alpha$-tubulin (n=3).

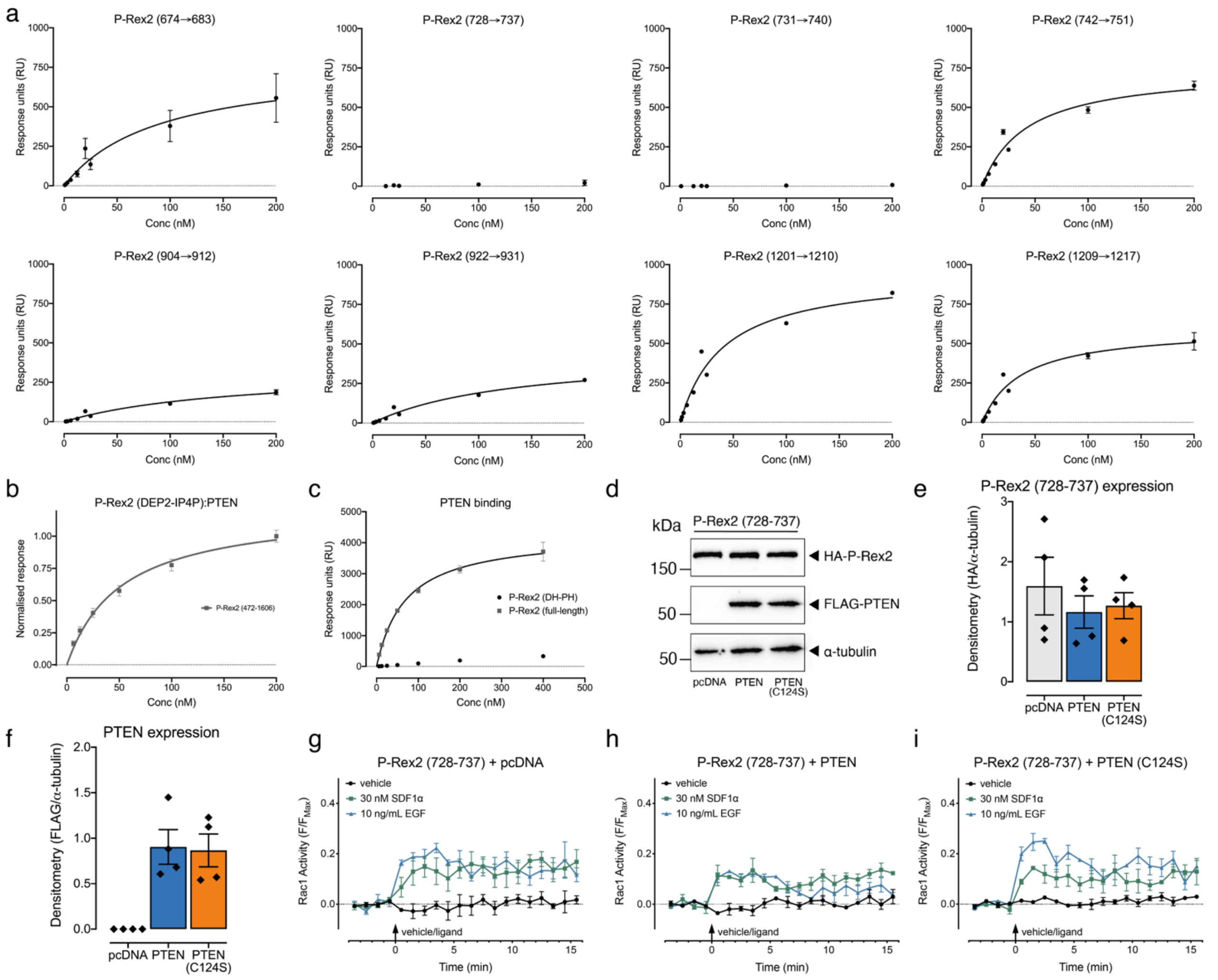


**Fig. S4. Characterisation of the PTEN:P-Rex2 interaction.** **a**, SPR steady-state binding curves of indicated P-Rex2 sequence swap mutants binding to PTEN. **b**, SPR steady-state response curves of PTEN binding by P-Rex2 (472-1606). **c**, SPR steady-state response curves of PTEN binding by P-Rex2 (full-length) compared to the isolated P-Rex2 (DH-PH) domains. **d**, Representative western blot showing HA-P-Rex2 and FLAG-PTEN (WT or C124S) expression following transfection in HEK293 cells. **e**, Quantification of HA-P-Rex2 (728-737) and **f**, FLAG-PTEN (WT or C124S) expression levels relative to α-tubulin (n=4). **g-i**, Time-course of Rac1 activation in HEK293 cells transiently expressing P-Rex2 (728-737) with pcDNA, PTEN, or PTEN (C124S).

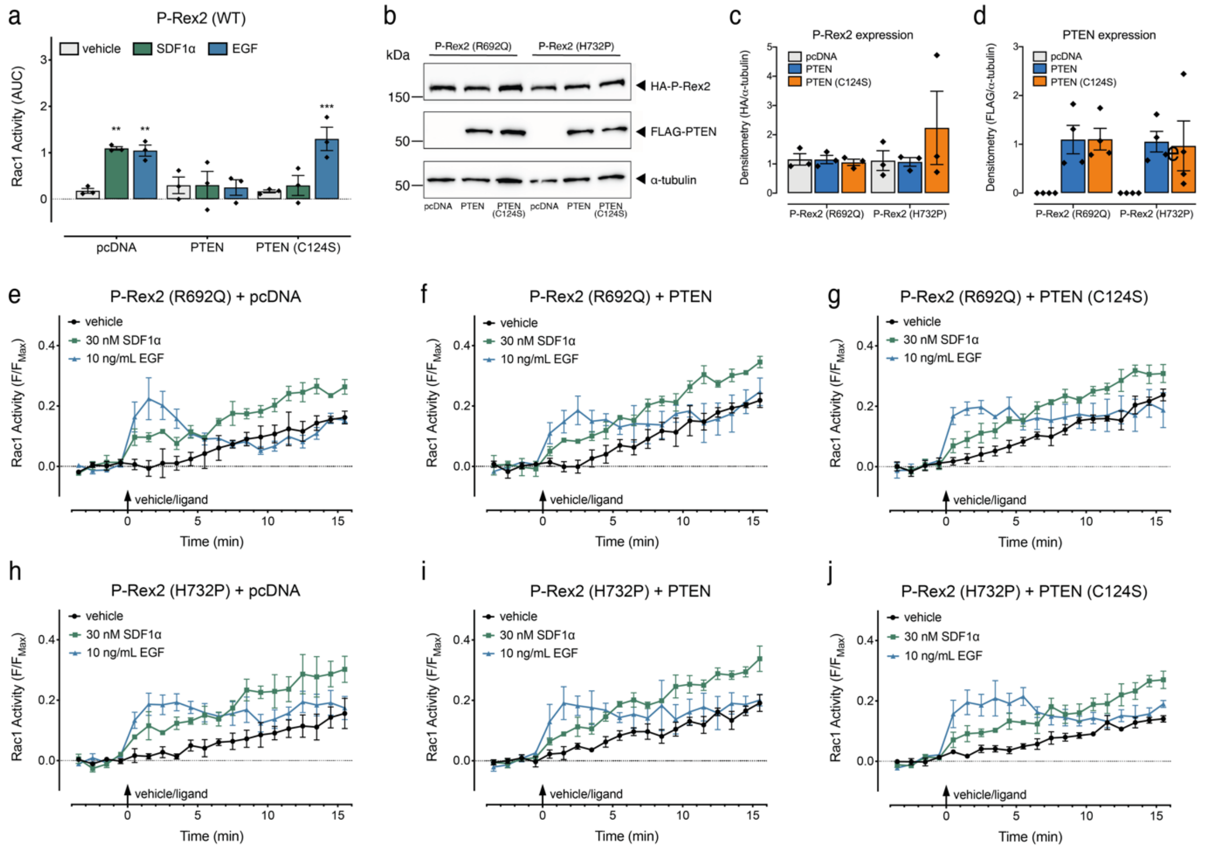


**Fig. S5. Characterisation of P-Rex2 PDZ2 cancer-associated somatic mutants. a**, Rac1 activity upon transient expression of P-Rex2 (WT) with pcDNA, PTEN, or PTEN (C124S) in HEK293 cells. AUC, the area under the curve calculated from **e-j**. Scatter plots show individual data points, symbols/bars represent means, and error bars indicate S.E.M. ** p<0.01 and *** p<0.001 versus vehicle control, two-way ANOVA with Dunnett's multiple comparison test. **b**, Representative western blot showing indicated HA-P-Rex2 somatic mutant and FLAG-PTEN (WT or C124S) expression following transfection in HEK293 cells. **c**, Quantification of HA (P-Rex2 somatic mutants) and **d**, FLAG (PTEN WT or C124S) expression relative to α-tubulin (n=3). **e-j**, Time-course of Rac1 activation upon expression of indicated P-Rex2 mutants with pcDNA, PTEN, or PTEN (C124S).

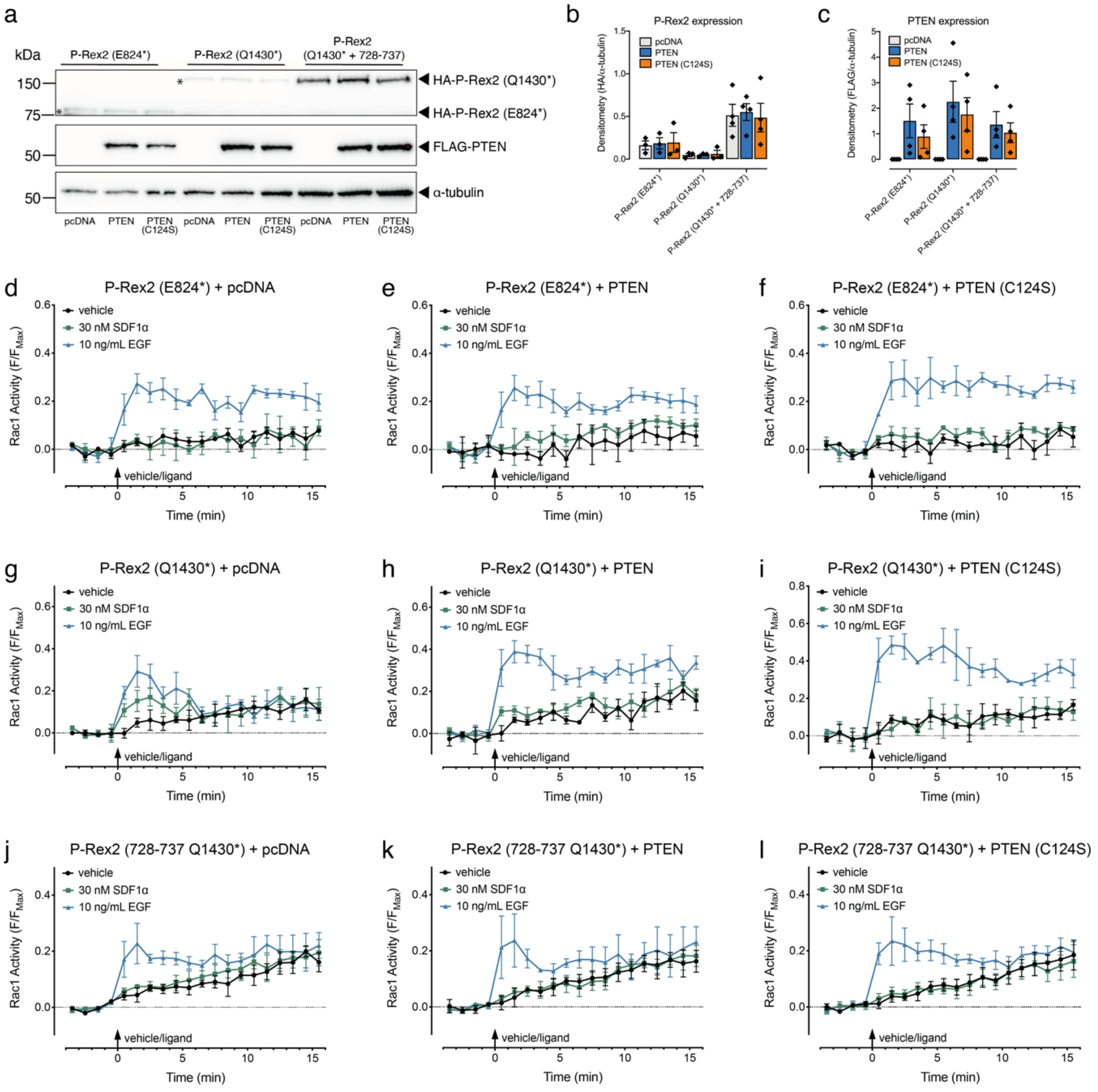


**Fig. S6. Characterisation of truncated P-Rex2 somatic cancer-associated mutants. a**, Representative western blot showing indicated HA-P-Rex2 somatic mutant and FLAG-PTEN (WT or C124S) expression following transfection in HEK293 cells. * indicates truncated P-Rex2 location. **b**, Quantification (from **a**) of HA (P-Rex2 somatic mutants) and **c**, FLAG (PTEN WT or C124S) expression relative to α-tubulin (n=4). **d-l**, Time-course of Rac1 activation upon expression of indicated P-Rex2 mutants with pcDNA, PTEN, or PTEN (C124S).
